# Computational reconstruction of the complete Piezo1 structure reveals a unique footprint and specific lipid interactions

**DOI:** 10.1101/783753

**Authors:** Jiehan Chong, Dario De Vecchis, Adam J Hyman, Oleksandr V Povstyan, Jian Shi, David J Beech, Antreas C Kalli

**Affiliations:** School of Medicine, University of Leeds, Leeds LS2 9JT, UK; Astbury Centre for Structural Molecular Biology, University of Leeds, Leeds, United Kingdom LS2 9JT

## Abstract

Piezo1 is a critical mechanical sensor in many cells. It is activated by mechanical force thus allowing cells to sense the physical environment and respond to stress. Structural data have suggested that Piezo1 has a curved shape. Here, we use computational approaches to model, for the first time, the 3D structure of the full-length Piezo1 in an asymmetric membrane. A number of novel insights emerge: (i) Piezo1 creates a dome in the membrane with a trilobed topology that extends beyond the radius of the protein, (ii) Piezo1 changes the lipid environment in its vicinity via specific interactions with cholesterol and PIP_2_ molecules, (iii) changes in cholesterol concentration that change the membrane stiffness result in changes in the depth of the dome created by Piezo1, and iv) modelling of the N-terminal region that is missing from current structures modifies Piezo1 membrane footprint, suggesting the importance of this region in Piezo1 function.

## INTRODUCTION

Piezo1 is a mechanosensitive ion channel found in many tissues. It plays key roles in the circulation (Choi et al., 2019; Li et al., 2014; Nonomura et al., 2018; Ranade et al., 2014; Retailleau et al., 2015; Rode et al., 2017), kidneys (Martins et al., 2016), red blood cells (Andolfo et al., 2013; Cahalan et al., 2015), and the central nervous system (Segel et al., 2019), acting in the development and maintenance of the mature organism, and ageing. Pathological mutations of Piezo1 lead to dehydrated hereditary stomatocytosis through gain of function (Albuisson et al., 2013; Bae et al., 2013), and generalized lymphatic dysplasia where there is loss of function (Fotiou et al., 2015; Lukacs et al., 2015).

Piezo1 is inherently mechanosensitive (Syeda et al., 2016; Wu et al., 2017b, 2016), but how it senses force remains largely unknown. Piezo1 is a non-selective cationic channel, with single channel conductance of ∼29 pS for mouse Piezo1 (mPiezo1) (Wu et al., 2017a). Structures of mPiezo1, excluding the N-terminal and parts of its cytosolic region, have been resolved by cryo-electron microscopy (cryo-EM) to resolutions of 3.7 – 4.8 Å (Ge et al., 2015; Guo and MacKinnon, 2017; Saotome et al., 2018; Zhao et al., 2018). These structures showed that Piezo1 adopts a trimeric configuration with a triskelion shape (Ge et al., 2015; Guo and MacKinnon, 2017; Saotome et al., 2018; Zhao et al., 2018). The C-termini of Piezo1 converge to form a central ion pore, while the N-termini extend from the central axis, forming “blades” which spiral in plane with the cell membrane as they radiate outward, as well as curving towards the extracellular surface (Figure 1). On the intracellular surface, a helical beam connects the peripheral regions of the blades to the central regions (Ge et al., 2015). So far, no structure for full-length Piezo1 has been published, thus limiting our ability to use structural data in order to understand Piezo1 function.

**Figure 1:**
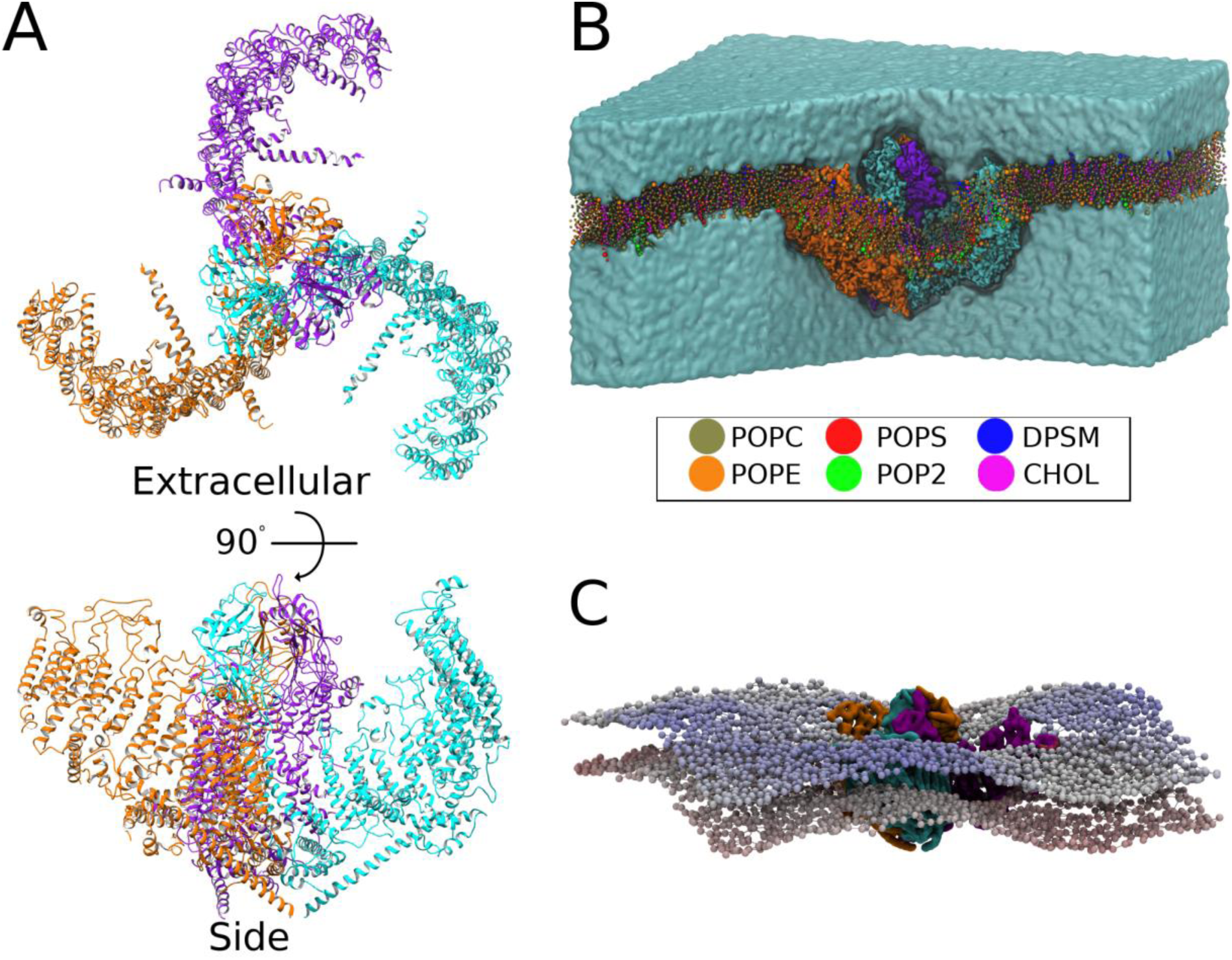
Piezo1 generates a complex unique membrane footprint. **(A)** Piezo1_trunc_ model, shown from an extracellular and side view. Piezo1 chains are shown in cyan, orange, and purple. **(B)** Snapshot from the end of the CHOL20_trunc_ simulation. Some of the solvent and bilayer were removed to illustrate the curvature of the bilayer induced by the transmembrane regions of Piezo1. Legend indicates colors for phosphatidylcholine (POPC), phosphatidylethanolamine (POPE), phosphatidylserine (POPS), phosphatidyl-4,5-biphosphate (PIP_2_), sphingomyelin (SM), and cholesterol (CHOL). **(C)** A snapshot from one of our CHOL20_trunc_ simulation. The Piezo1_trunc_ backbone is displayed in surface representation. Phosphate beads are colored by their z-coordinates, from red at the bottom to blue at the top, illustrating crests and troughs in the bilayer.

Membrane lipids have been shown to regulate ion channel function (Rosenhouse-Dantsker et al., 2012). In particular, cholesterol has the potential to affect the mechanogating of Piezo1, due to its effect on membrane stiffness (Needham and Nunn, 1990). The effect of cholesterol on other ion channel families is varied. Cholesterol addition has been reported to increase the activity of Orai1 (Derler et al., 2016), Kv1.3 (Hajdú et al., 2003), Kir2 (Romanenko et al., 2002) and the sodium inward current (Wu et al., 1995). In other studies, cholesterol depletion reduces the activity of ENaC (Balut et al., 2005) and TRPC (Bergdahl et al., 2003) channels. In a recent preprint study, lateral organization of cholesterol in the membrane is reported to regulate Piezo1 activity (Ridone et al., 2019). Further indirect links between cholesterol and Piezo1 function exist through STOML3, which tunes Piezo1 activity (Poole et al., 2014) as well as membrane stiffness (Qi et al., 2015). Cholesterol has previously been shown to regulate the activity of Ca^2+^-permeable stretch-activated channels (Chubinskiy-Nadezhdin et al., 2011; Morachevskaya et al., 2007), but it is not known whether these channels were Piezo1. Additionally, PIP_2_ has been suggested to regulate Piezo1 function (Borbiro et al., 2015). Recent studies have also shown that molecules e.g. fatty acids that alter the membrane structural properties regulate Piezo1 activity (Conrard et al., 2018; Romero et al., 2019).

A recent study has shown that the unique curved shape of Piezo1 induces a dome in the membrane, possibly creating an energy store which may provide a basis for mechanogating (Guo and MacKinnon, 2017). Using mechanical calculations in which the dome was modeled as hemispherical it was suggested that membrane tension would flatten the dome, providing energy which could open the Piezo1 channel. In addition to the membrane region curved by direct contact with Piezo1, it was also suggested that the local Piezo1-induced curvature generates a wider membrane footprint, which is hypothesized to amplify the sensitivity of Piezo1 to mechanical force (Haselwandter and MacKinnon, 2018). The sensitivity of Piezo1 is tuned by membrane tension and stiffness, which may in turn be influenced by varying membrane lipid composition (Lewis and Grandl, 2015; Qi et al., 2015; Romero et al., 2019). A ‘force from lipids’ model of mechanical activation has been proposed, in which nothing is required for Piezo1 mechanosensation other than the cell membrane and Piezo1 itself (Lewis and Grandl, 2015; Qi et al., 2015). Other proposed mechanisms for Piezo1 force sensing include shear force deflection of the extracellular cap domain, membrane thinning in response to stretch, and changes to membrane lipid composition (Wu et al., 2017a). In addition to mechanical force, Piezo1 is also selectively activated by the chemical agonist, Yoda1 (Syeda et al., 2015).

Despite the recent functional and structural evidence about Piezo1 our understanding of its footprint in the membrane and how lipids may regulate its function is still limited. This is partly because the structure of the full-length Piezo1 is not available, and partly due to the fact that critical details about the topology of Piezo1 footprint in the membrane and about its interaction with its lipid environment are currently missing. In this study, we used the available structural data about Piezo1 to provide a three-dimensional (3D) full-length Piezo1 structural model. We have used this model to perform molecular dynamics (MD) simulations and combined this with lab-based methodologies to show, for the first time, the unique topology of the Piezo1 membrane footprint and how this is regulated by cholesterol concentration. Moreover, this study demonstrates the critical role of the N-terminal region of Piezo1 in regulating its local membrane environment and suggests interaction sites for anionic PIP_2_ lipids, supporting their proposed functional role (Borbiro et al., 2015).

## RESULTS

### Piezo1 generates a trilobular dome and extensive penumbra

A structural model of the truncated Piezo1 (Piezo1trunc) was generated by adding the missing loops to the published structural data of the mouse protein (PDB: 6B3R; see Methods and Figure 1A). We simulated Piezo1_trunc_ using coarse-grained molecular dynamics (CG-MD) simulations, in a complex lipid bilayer containing 20% cholesterol and a full complement of phospholipids and sphingomyelin (see Supplementary Table 1). This lipid mixture mimics the physiological endothelial membrane (Medow et al., 1989; Murphy et al., 1992; Takamura et al., 1990). Although the Piezo1_trunc_ model lacks the 3 distal 4-helical bundles in its blades, our simulations suggest that it changes the curvature of the membrane in its immediate vicinity, creating a stable membrane dome (Figure 1B). This local deformation results in a wider membrane deformation with a complex three-dimensional topology, which extends beyond the vicinity of Piezo1_trunc_. This footprint creates elevated crests in the bilayer which radiate outward from the blade tips of Piezo1_trunc_ (Figure 1C, in blue), whilst depressed valleys radiate from the regions between the arms (Figure 1C, in white). Therefore, despite the absence of some of its N-terminal region Piezo1_trunc_ creates a unique trilobed membrane topology that extends outside the radius of Piezo1_trunc_, that could potentially regulate its function and interactions.

The precise function and structure of the Piezo1 N-terminal residues 1-576 remains uncertain (Ge et al., 2015; Guo and MacKinnon, 2017; Zhao et al., 2018). Given that the presence of the missing N-terminal region will extend the Piezo1 transmembrane region as suggested by a recent structure of the homologous Piezo2 protein (PDB: 6KG7) we hypothesized that the structurally unresolved N-terminal residues might impact on the dimensions and dynamics of the Piezo1 footprint. To address this question, we built a full-length structural model for Piezo1. Using secondary structure prediction tools (see Methods) we generated a topology for the N-terminal bundles which extends the blades as previous suggested (Guo and MacKinnon, 2017). To build our model, we took advantage of the fact that the three 4-helical bundles that are missing have the same structure/topology with the adjacent group of three 4-helical bundles in Piezo1 blades i.e. residues 577-1129 (bundles 4-5-6) of the resolved Piezo1 structure (PDB: 6B3R) to model the missing N-terminal residues. This model completes the Piezo1 structure by including cytoplasmic residues missing from the published structure (see Methods). The resulting model for the full-length Piezo1 (Piezo1_full_) is shown in Figure 2A, providing the first 3D model of the full-length Piezo1.

**Figure 2:**
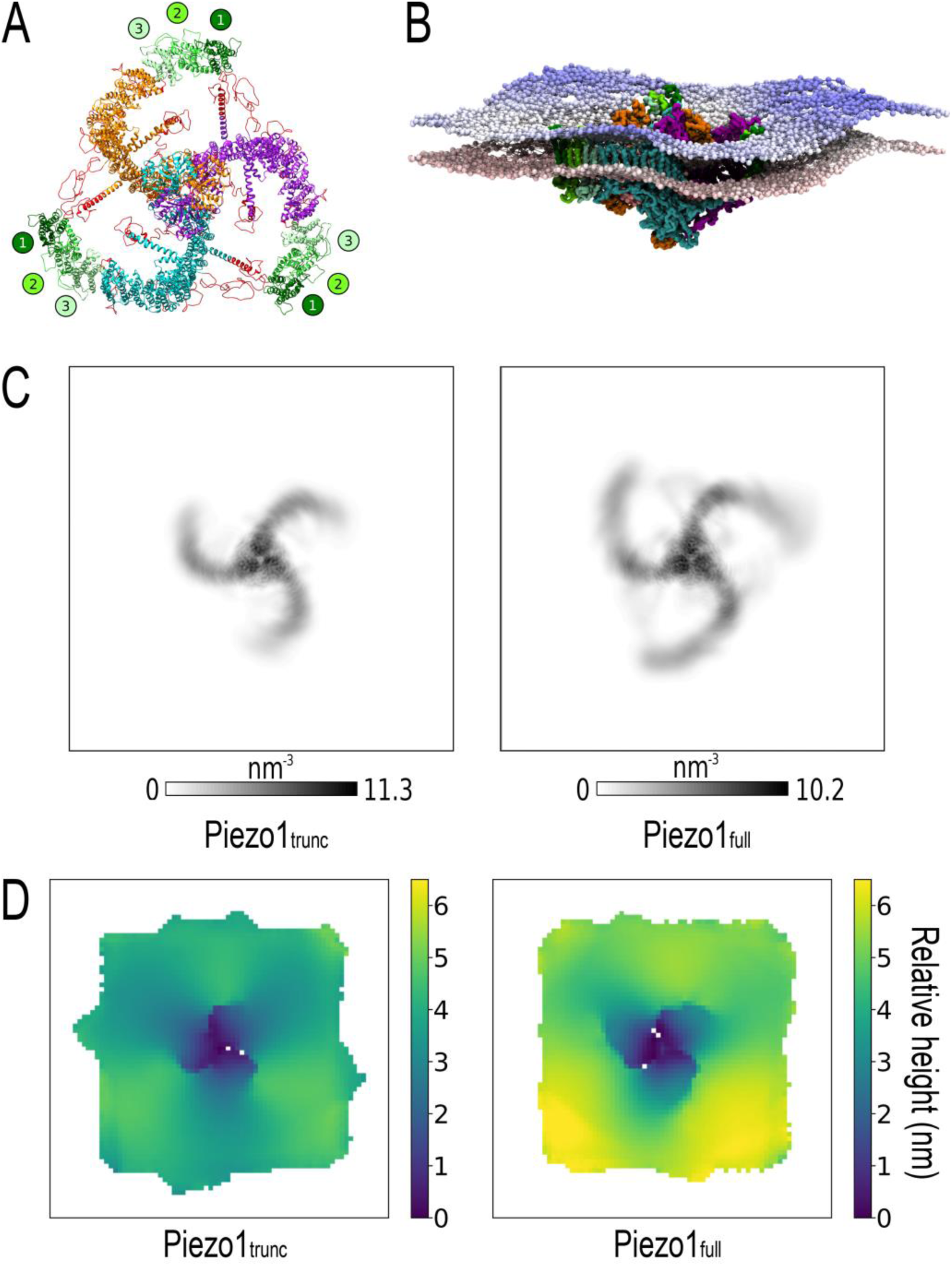
The full-length Piezo1 model has flexible distal N-terminal blades and enhances the footprint produced by the truncated mPiezo1 structure. **A)** Extracellular view of the full-length mPiezo1 model. Chains are colored as in Figure 1A. The modeled transmembrane N-terminal bundles (residues 1-576) are numbered and colored from dark to light green. Other residues not resolved in the cryo-EM structure are colored red. **B)** Final snapshot from one of the CHOL20_full_ simulations. The Piezo1_full_ backbone particles are displayed in surface representation. Phosphate beads are colored according to their z coordinates from red at the bottom to blue at the top, illustrating the complex topology of the bilayer footprint. **C)** 2-dimensional protein density of Piezo1_trunc_ and Piezo1_full_ during CHOL20_trunc_ and CHOL20_full_ simulations respectively. Coordinates are fitted to the pore region (residues 2105-2547). **D)** Height maps of CG phosphate beads corresponding to the extracellular leaflet in CHOL20_trunc_ (left) and CHOL20_full_ (right) simulations, averaged across all repeats.

In our Piezo1_full_ model each blade from a single chain is composed of 36 α-helices that form a total of 9 4-helix bundles as previously suggested (Guo and MacKinnon, 2017) (Supplementary Figure 1). The Piezo1_full_ model also comprises cytoplasmic residues not resolved by cryo-EM (Guo and MacKinnon, 2017) (Figure 2A, red ribbons).The assignment of loops to extracellular or intracellular positions agrees with previous experimental results, which used a combination of myc tagging and phosphorylation sites to distinguish extracellular and intracellular regions of Piezo1 (Coste et al., 2015) (Supplementary Figure 1C).

In order to assess how the full length Piezo1 will alter the membrane footprint, CG-MD simulations with the Piezo1_full_ model inserted in a bilayer with identical lipid composition to CHOL20_trunc_ (Supplementary Table 1) were performed. In all five repeat simulations performed, the modeled N-terminal bundles remained embedded in the lipid bilayer (Figure 2B). To inspect the extent of blade movement over the course of the simulation, we analyzed the 2-dimensional protein density for both Piezo1_trunc_ and Piezo1_full_ structural models (Figure 2C). We found that the protein density decreases in magnitude and becomes more diffuse as we move outward from the proximal C-terminal pore region towards the distal N-terminal blades. This effect is amplified in Piezo1_full_, suggesting that the modelled N-terminal residues impart a wider range of movement to the blades, especially to the N-terminal region. This might explain why structural techniques were unable to resolve the structure of the N-terminal region of Piezo1.

To analyze the topology of the Piezo1 footprint, we used a script that we developed (see Methods) to generate an average height map of CG phosphate beads in each leaflet (Figure 2D). Surprisingly, this analysis reveals a footprint that is not uniformly circular, but trilobate as seen for the Piezo1_trunc_. In comparison to Piezo1_trunc_, Piezo1_full_ produces a more pronounced footprint, with maximum depth exceeding 6 nm. In addition, the Piezo1_full_ model produces a footprint with wider crests, and narrower valleys. This result confirms the capacity of Piezo1 to indent the membrane and establishes a convoluted topology for the wider Piezo1 membrane footprint, to which the blades make an important contribution. Moreover, the deepest point of the dome is on the pore region.

### The depth of the Piezo1 footprint varies with cholesterol concentration

It is suggested that the dimensions of the Piezo1 footprint may affect the sensitivity of Piezo1 to mechanical stress (Haselwandter and MacKinnon, 2018). Changes in the percentage of cholesterol are expected to affect the properties of the bilayer and in particular its curvature. To determine whether membrane cholesterol concentration influences the dimensions of the Piezo1 footprint, we performed further CG-MD simulations of Piezo1_trunc_ in a range of complex bilayers with cholesterol concentrations of 0%, 5%, 10%, 30%, and 40%, and Piezo1_full_ in cholesterol concentrations of 0%, 10%, 20%, and 30% (Supplementary Table 1). For each cholesterol concentration, the depth of the Piezo1 footprint was analyzed using our script as above (see Methods).

In the Piezo1_trunc_ system, the deepest footprint is observed at 0% cholesterol, with a secondary peak at 30% cholesterol. The footprint is most shallow in between these peaks, at 10% cholesterol. In the Piezo1_full_ system, the same reduction is observed between 0% and 10% but unlike the Piezo1_trunc_ system the depth of the dome remains almost the same when cholesterol concentration increases to 20% (Figure 3). Similar to the Piezo1_trunc_ system the depth of the dome increases at 30% cholesterol. These data show that depth of the Piezo1 footprint varies non-linearly with cholesterol concentration, following an inverted unimodal relationship with a minimum depth at 10% cholesterol for the Piezo1_trunc_ and 10 – 20% cholesterol for Piezo1_full_.

**Figure 3:**
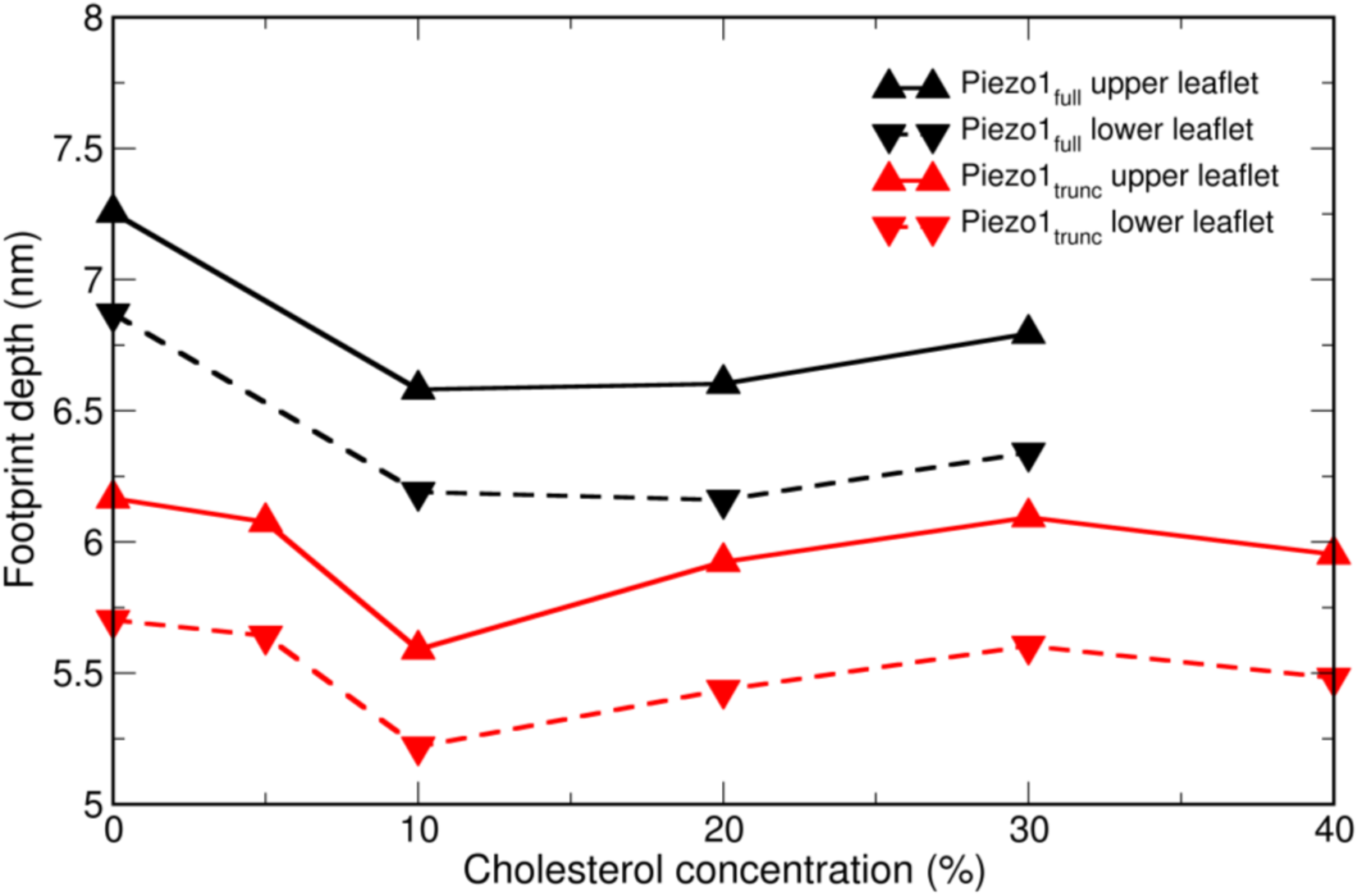
The depth of the Piezo1 membrane dome varies with cholesterol concentration. Average depth of the Piezo1 membrane dome in the CG-MD simulation with different membrane cholesterol concentrations. Systems and leaflets are indicated in the figure legend.

### Changes of cholesterol concentration suppress Piezo1-dependent Ca^2+^ entry

Ca^2+^ influx has been found to be an endogenous shear stress response of endothelial cells that is dependent on Piezo1 (Li et al., 2014). Our simulations above suggest that cholesterol concentration affects the depth of the Piezo1 footprint, possibly regulating its function (Haselwandter and MacKinnon, 2018). To determine the effects of cholesterol we have performed experiments with Yoda1 which is a Piezo1 channel agonist (Syeda et al., 2015). To confirm an effect of Yoda1 on Piezo1 in human umbilical vein endothelial cells (HUVECs), we tested the effect of Yoda1 on Ca_2+_ influx, with and without the presence of Piezo1 siRNA. At a sub-maximal concentration, Yoda1 produced a large Ca_2+_ influx reaching a maximum at 90 seconds. This effect is suppressed by Piezo1 siRNA (Supplementary Figure 2).

Treating HUVECs with exogenous cholesterol reduced Ca_2+_ influx in response to Yoda1 (Figure 4A, left). The IC_50_ of cholesterol was 0.04 mM (Figure 4A, right). However, due to its limited solubility, higher concentrations of cholesterol were not tested which may have given a more complete block of Yoda1 responses.

**Figure 4:**
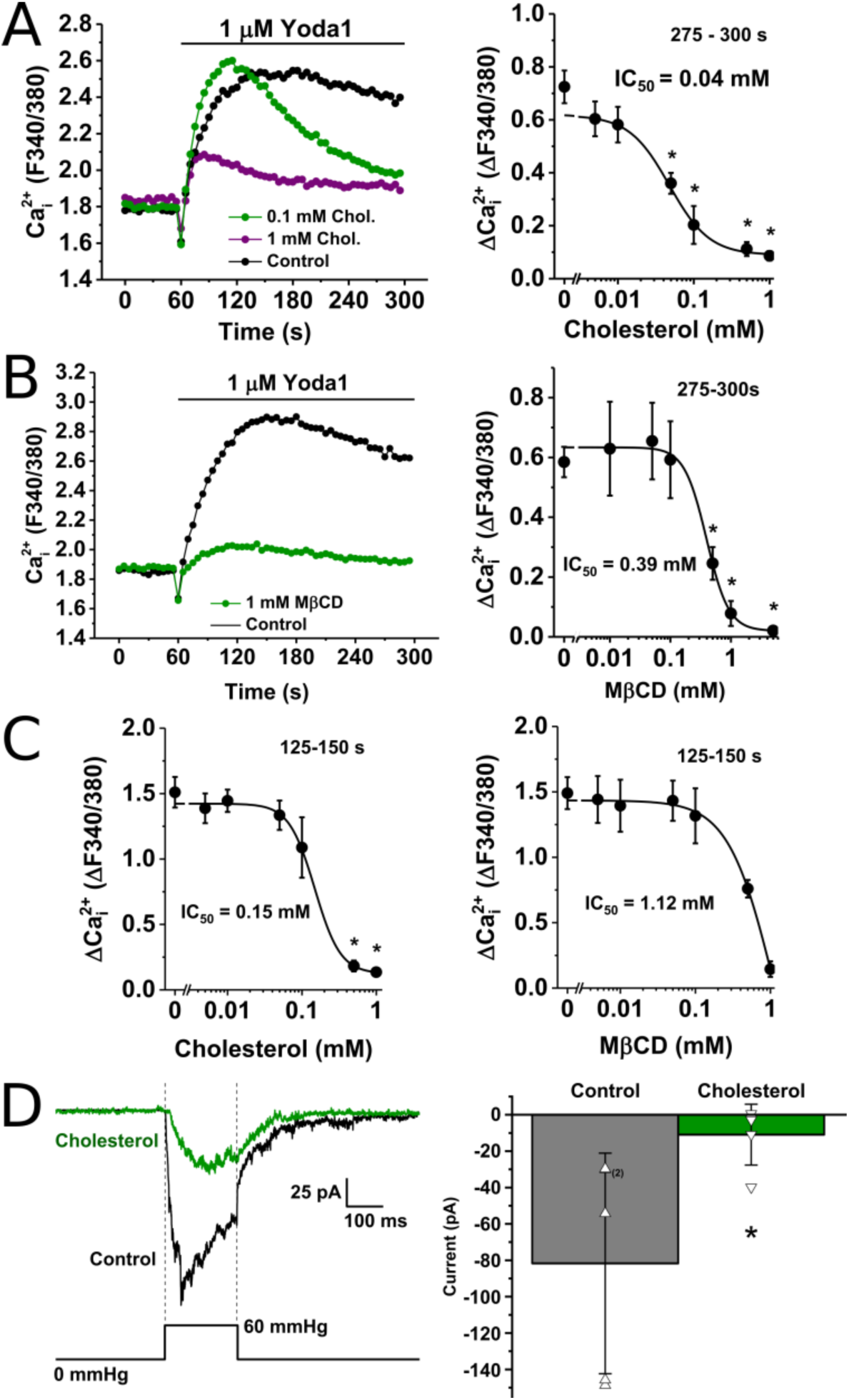
Cholesterol modulates Piezo1 responses to Yoda1 and mechanical activation. **(A)** Left, an example 96-well plate fura-2 measurement of the change in intracellular Ca^2+^ concentration evoked by 1 µM Yoda1 in HUVECs pre-treated with 0 (control), 0.1, and 1 mM cholesterol. Right, mean data for the average amplitude (at 275-300 seconds) of HUVEC responses to Yoda1 with varying doses of cholesterol (0.005-1 mM) displayed as a Hill Equation indicating the 50 % inhibitory effect (IC50) at 0.04 mM. n/N=4/4. (**B)** Left, an example 96-well plate fura-2 measurement of the change in intracellular Ca^2+^ concentration evoked by 1 µM Yoda1 in HUVECs pre-treated with 0 (control) and 1 mM MβCD. Right, mean data for the average amplitude (at 275-300 seconds) of HUVEC responses to Yoda1 with varying doses of MβCD (5-0.01 mM) displayed as a Hill Equation indicating the 50 % inhibitory effect (IC50) at 0.39 mM. n/N=4/4. **(C)** Left, mean data of peak amplitudes, taken between 125 – 250 seconds, of the response of Human Embryonic Kidney cells conditionally expressing Piezo1 (P1 HEK T-Rex cells) to Yoda1 with varying doses of cholesterol (0.005 – 1 mM). n/N=4/4. Right, mean data of peak amplitudes, taken between 125 – 250 seconds, of P1 HEK T-Rex cell responses to Yoda1 with varying doses of MβCD (0.005 – 1mM). n/N=4/4. * = P < 0.05. **(D)** Recordings were made using outside-out patch configuration applied to mouse P1 T-REx-293 cells. Currents were evoked by 200-ms positive pressure steps of 60 mmHg applied to the patch pipette. The holding potential was −80 mV. Left, example current traces recorded from two different patches pre-incubated with control solution (black trace) or solution containing 0.1 mM cholesterol applied for 30 min at room temperature prior to recording (green trace). Right, mean ± s.d. of peak current for experiments of the type shown on the left, with all original data points superimposed (n=5 for each group).

As the results above suggest that addition of cholesterol has an inhibitory effect on Piezo1 activity, the importance of endogenous cholesterol in HUVEC membranes was investigated. This was achieved by chemically depleting membrane cholesterol using methyl-β-cyclodextrin (MβCD), a cholesterol-binding compound. MβCD reduced the amplitude of Yoda1 responses in HUVECs (Figure 4B, left). High concentrations of MβCD reduced the amplitude of Yoda1 responses in HUVECs with an IC_50_ of 0.39 mM (Figure 4B, right). MβCD concentrations in excess of 1mM reduced cell attachment, compromising the quality of Ca_2+_ measurement data. Negative controls treated with α cyclodextrin, a compound with similar structure to MβCD but which does not bind cholesterol, showed no change in Yoda1 responses (Supplementary Figure 3).

To clarify whether the effect of cholesterol on Piezo1 is an inherent property of the protein, or due to the specific HUVEC environment, the effect of cholesterol on Yoda1 responses was tested in an overexpression system. In T-Rex HEK 293 cells with conditional human Piezo1 expression (P1 HEK T-Rex), the addition of exogenous cholesterol, and removal of cholesterol using MβCD, reduces the amplitude of Yoda1 responses in a manner echoing our observations in HUVECs (Figure 4C).

As Yoda1 activation of Piezo1 is modulated by cholesterol, we sought to determine if mechanical activation is similarly affected by cholesterol. In HEK 293 cells stably expressing mouse Piezo1, to exactly match the model of mouse Piezo1, the addition of exogenous cholesterol inhibits mechanically activated currents (Figure 4D).

Together, these results suggest a bimodal relationship between cholesterol concentration and Piezo1 activity, where either addition or removal of cholesterol leads to a reduction in Piezo1 activity.

### Piezo1 interacts with cholesterol in a model bilayer

To examine the interaction between cholesterol and Piezo1 in more detail, cholesterol density and protein-cholesterol contacts were analyzed in our CG-MD simulations of Piezo1_trunc_ and Piezo1_full_. In our simulations, cholesterol clusters alongside structural regions, e.g. between the helices TM30 and TM31, TM22 and TM27, and adjacent to TM15 (Figure 5A, B). Using 60% of maximum contacts as a cut-off, 64 Piezo1_trunc_ (Supplementary Table 2) and 62 Piezo1_full_ (Supplementary Table 3) residues have significant contacts with cholesterol in 1 or more simulations. This regional clustering is observed across all cholesterol concentrations tested (Supplementary Figure 4). The N-terminal region of Piezo1, which is not present in the published structures, also makes significant interactions with cholesterol (Supplementary Table 3).

**Figure 5:**
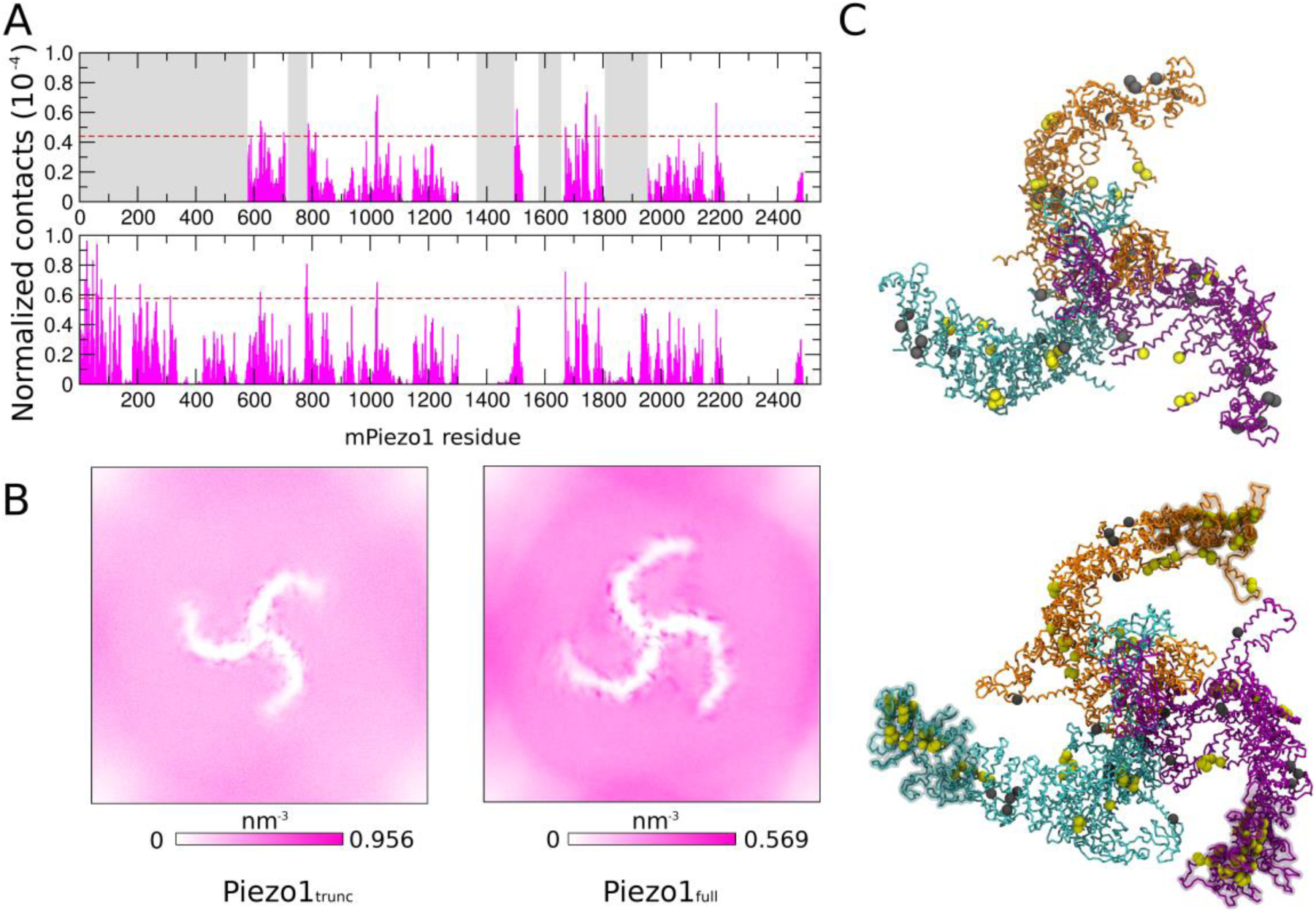
Piezo1 interactions with cholesterol in a complex model bilayer. **(A)** Normalized contacts between cholesterol and Piezo1_trunc_ (top) and Piezo1_full_ (bottom) in the model bilayer containing 20% cholesterol. Normalization was done by dividing the number of cholesterol contacts of each residue by the number of cholesterol molecules and total number of frames analyzed. The greyed-out regions of the Piezo1_trunc_ plot represent residues absent in Piezo1_trunc_. Red dotted lines represent the cutoff of 60% of the contacts in each simulation. **(B)** Two-dimensional density maps of cholesterol over 5 repeat simulations of CHOL20_trunc_ and CHOL20_full_, showing clustering of cholesterol along the Piezo1 blades. **(C)** Snapshots of Piezo1_trunc_ (top) and Piezo1_full_ (bottom), with dynamic bonds between backbone residues colored as in figure 1A. Cholesterol-interacting residues are displayed as yellow or grey spheres. Residues in yellow made significant interactions with cholesterol. Residues in grey spheres both made significant interactions with cholesterol and are part of CRAC/CARC motifs. The modelled N-terminal regions of Piezo1_full_ are displayed as transparent surfaces

CRAC and CARC consensus motifs are noted cholesterol-binding domains (Fantini and Barrantes, 2013). To determine the relevance of these motifs to Piezo1-cholesterol interactions, we searched for these motifs in the Piezo1 sequence (Supplementary Table 4). 19 CRAC and 40 CARC motifs were identified. Of these, 4 CRAC and 10 CARC sequences overlap with residues with significant cholesterol interactions in either Piezo1_full_ or Piezo1_trunc_ (Figure 5C, top, Supplementary Table 4).

### Predicted specific interaction of full-length Piezo1 with PIP_2_

Lipid species other than cholesterol might also impact on Piezo1. For instance, phosphatidyl-4,5-biphosphate (PIP_2_), a minority membrane lipid, is an important regulator of ion channels (Huang, 2007; Suh and Hille, 2008), including Piezo1 (Borbiro et al., 2015). The use of a complex asymmetric bilayer (Supplementary Table 1) in our simulations allows us to also examine the interactions of Piezo1 with other lipids. Interestingly, in our analysis, PIP_2_ interacts strongly with specific Piezo1 residues (Figure 6A). Across all simulations, there were 33 Piezo1_trunc_ residues (Supplementary Table 5) and 43 Piezo1_full_ residues (Supplementary Table 6) with contacts above the cutoff of 70% of the contacts in one or more systems. Of these, 17 are present in all Piezo1_trunc_ systems (Supplementary Table 5). 23 residues are over cutoff in all Piezo1_full_ systems (Supplementary Table 6). These include missing loops and bundles that have been modelled in Piezo1_full_. 10 residues are above cutoff in all systems for both models (R629, K630, R633, R844, R846, R1023, R1024, R1025, K1201, R1204). In simulations of both Piezo1_trunc_ and Piezo1_full_, PIP_2_ lipids form an anionic annulus around Piezo1 in the inner leaflet (Figure 6B). Similar specific PIP_2_ clustering (Supplementary Figure 5) and annulus formation (Supplementary Figure 6) is observed across all cholesterol concentrations tested.

**Figure 6:**
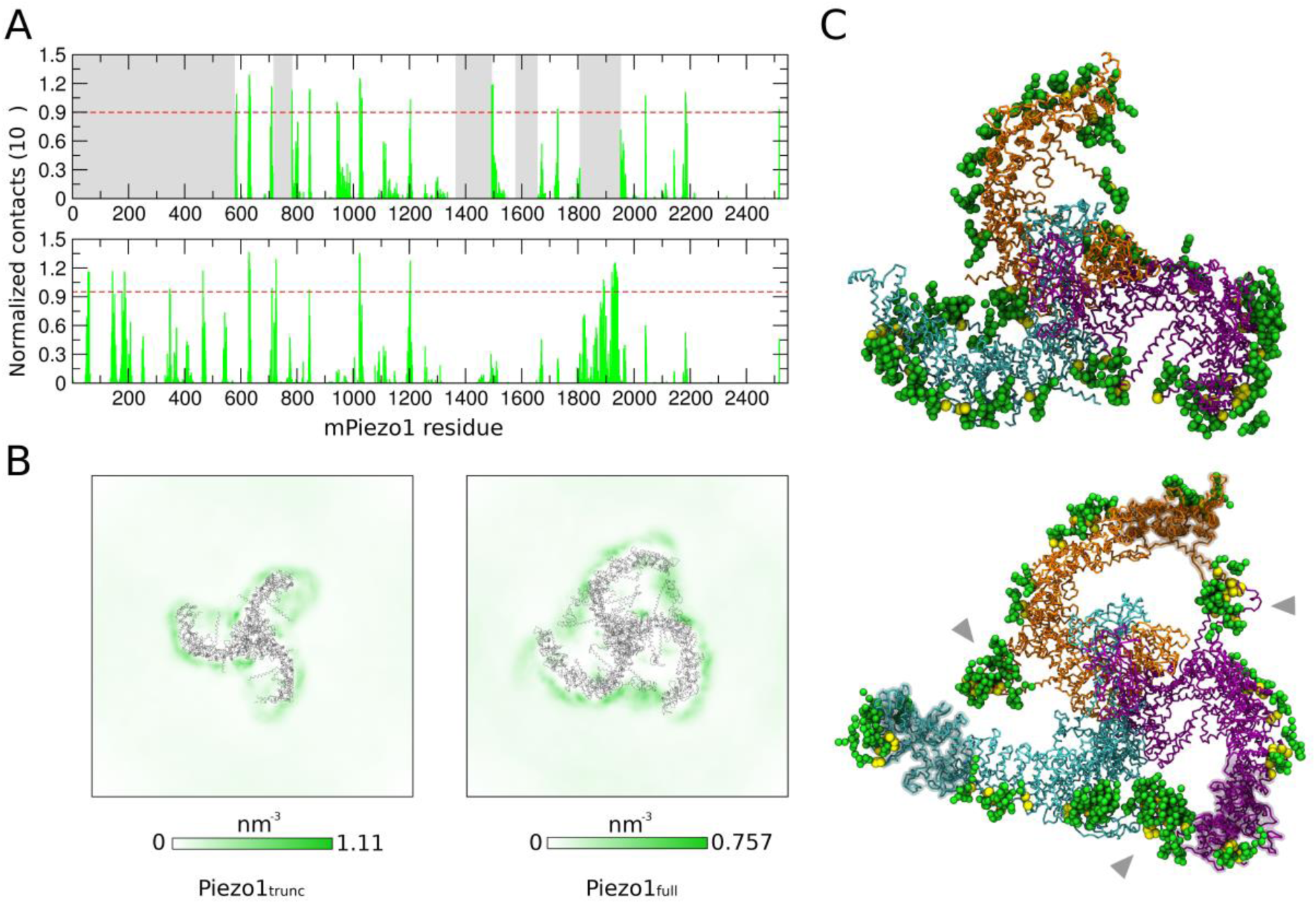
Piezo1 interactions with PIP_2_ lipids. **(A)** Normalized number of contacts between PIP_2_ and Piezo1_trunc_ (top) and Piezo1_full_ (bottom) in the CHOL20_trunc_ and CHOL20_full_. Greyed out regions represent residues absent in Piezo1_trunc_. Red dotted lines represent the cutoff of 70% of the contacts in each simulation. **(B)** 2D density maps of PIP_2_ lipids in CHOL20_trunc_ (left) and CHOL20_full_ (right). **(C)** Snapshots from one of our CHOL20_trunc_ (top) or CHOL20_full_ (bottom) simulations. The Piezo1 backbone are shown in the same colors as Figure 1A. Residues with PIP_2_ contacts exceeding the cutoff of 70% maximum contacts are displayed as yellow spheres. PIP_2_ lipids with head groups within 10 Å of the Piezo1 residues that forms significant contacts with PIP_2_ are displayed as green spheres. Grey arrows point to PIP_2_ lipids clustering at the interface between adjacent Piezo1_full_ blades.

Interestingly, the PIP_2_ annulus around Piezo1_full_ differs significantly from that around Piezo1_trunc_. In Piezo1_full_ the PIP_2_ clustering occurs at the interface between the helices that lie in the plane of the inner leaflet, and the convex surface of the adjacent blade (Figure 6C, bottom, grey arrows). There are regions of low PIP_2_ density between the concavity of each blade and the central core of Piezo1, which may be a consequence of PIP2 being drawn preferentially to the interfacial clusters (Figure 6B, right). This behavior of PIP_2_ around Piezo1_full_ is consistent across all cholesterol concentrations tested (Supplementary Figure 5) and suggest that the presence of the N-terminal region of Piezo1 changes this shape of the PIP_2_ annulus around Piezo1.

Contact analysis of POPC (Supplementary Figure 7), POPE (Supplementary Figure 8), POPS (Supplementary Figure 9), and sphingomyelin (SM) (Supplementary Figure 10) was performed, as well as density mapping of all lipid species simulated (Supplementary Figure 5). We did not observe specific clustering or annulus formation for any lipid species other than PIP_2_.

## DISCUSSION

In this study we model, for the first time, in molecular detail the Piezo1 full-length structure and its membrane footprint. Although it has previously been suggested that in the inactive state the Piezo1 N-terminal blades possibly flatten rather than continuing to curve (Guo and MacKinnon, 2017), our approach of using the existing transmembrane bundles as a template leads to a continuous curve in the modelled region. This result is supported by a recent cryo-EM structure for the homologous mechanosensor Piezo2, in which the N-termini also continue to curve (Wang et al., 2019). Comparison of our Piezo1_full_ model with the Piezo2 structure shows that the overall structures are very similar (Supplementary Figure 1E). The N-terminal region of the blades of the Piezo2 structure is more elevated relative to our Piezo1_full_ structure (Supplementary Figure 1F), however the Piezo1 blades are highly flexible in response to force (Lin et al., 2019), and the same is likely to be true of Piezo2. Therefore, differences between these static structures may not represent dynamic differences *in vivo*.

Our results suggest that Piezo1 footprint has a trilobed topology with a dome facing into the cytosolic region of Piezo1 that extends beyond the immediate vicinity of the protein. Uniquely, our results also show that Piezo1 does not only change the membrane curvature around the protein, but it also changes its local lipid environment by creating an anionic annulus in the inner leaflet and by strong interactions with cholesterol. Interestingly, the full length Piezo1 structure alters this membrane footprint and its relationship with cholesterol, by increasing the dome depth, by changing the profile of its peaks and troughs, and by modifying its interactions with cholesterol and PIP_2_. This hints that the N-terminal region plays a critical role in the topology of the membrane surrounding Piezo1. Our results also provide novel insights into the contribution of cholesterol to the depth of the dome created by Piezo1.

This unique Piezo1 footprint appears to be a long-range consequence of the immediate membrane indentation induced by Piezo1. A study by Guo and Mackinnon (Guo and MacKinnon, 2017) showed that the depth of the dome created by Piezo1 is ∼6 nm. This is consistent with the 6-7 nm seen in our simulations. We note that our results also suggest that the depth of the dome in the full length Piezo1 is larger compared to the truncated, highlighting the role of the N-terminal region of the blades in regulating the Piezo1 membrane deformation. A wider perturbation of the membrane surrounding Piezo1 has also been hypothesized (Haselwandter and MacKinnon, 2018) using mechanical calculation and approximating the dome as a hemispherical dome. Our study shows that the Piezo1 footprint is far larger than Piezo1 radius but for the first time suggests that it has a uniquely complex tri-lobed topology with regions of varying curvature. This observation suggests that Piezo1 may be not only a passive receptor of mechanical forces but could transmit physical effects at long range through the membrane. This idea of Piezo1 applying force to the membrane while the membrane applies force to the protein is supported by recent work from the MacKinnon and Scheuring groups (Lin et al., 2019). The topology may also amplify Piezo1 tension sensitivity as previously suggested (Guo and MacKinnon, 2017).

Membrane curvature is believed to be not just a consequence of cellular processes, but also a generator of downstream effects including membrane scission, fusion, protein sorting, and enzyme activation (Chang-Ileto et al., 2011; McMahon and Boucrot, 2015; McMahon and Gallop, 2005; Reynwar et al., 2007). The complex topology of the Piezo1 footprint, with regions of different height and curvature, raises intriguing possibilities regarding how Piezo1 could interact with membrane protein partners. Stable regions of differential curvature in the membrane might effectively function as coplanar compartments, with segregation of cellular processes by curvature. Membrane tension could be expected to alter the curvature difference between compartments, leading to alterations in protein aggregation (Reynwar et al., 2007) or lipid signalling (Chang-Ileto et al., 2011). Hence, the complex Piezo1 footprint offers an alternative pathway for Piezo1-dependent mechanotransduction, beyond its properties as an ion channel. This possibility could be investigated using CG-MD simulation on a larger scale, or experimentally with imaging techniques such as Förster Resonance Energy Transfer (FRET) microscopy or Stochastic Optical Reconstruction Microscopy (STORM).

The Piezo1_full_ model offers unique insight into Piezo1 structure. The high mobility of the N-terminal residues in our simulations (Figure 2) may explain why to date, it has been challenging to resolve the structure of this region by cryo-EM (Guo and MacKinnon, 2017; Saotome et al., 2018; Zhao et al., 2018). Comparison of the transmembrane region of the Piezo1_full_ model with the transmembrane region of the homologue Piezo2 demonstrates that the two proteins have a very similar shape.

Furthermore, compared to Piezo1_trunc_, Piezo1_full_ changes the effect on membrane curvature, modulates the effect of cholesterol on footprint depth, and alters predicted lipid interactions with PIP_2_ and cholesterol. These differences highlight the importance of the N-terminal region that is currently missing from the current structure. Extension of the Piezo1 blades increases the depth of the Piezo1_full_ footprint relative to Piezo1_trunc_ (Figures 2D and 3). This observation couples Piezo1 blade configuration to local membrane curvature and supports the idea that the blades create, as well as sense, membrane curvature. Additionally, the high degree of mobility at the N-terminal ends of the blades may supports a model in which Piezo1-induced curvature is dynamic and can change in response to force.

The Piezo1_full_ model also includes extended unstructured loops on the cytoplasmic region that are also missing from the published structure (Figure 2A, red ribbons). Despite their lack of secondary structure, these loops contain numerous charged residues, and may represent hotspots for cytoskeletal interactions or post-translational modification (e.g., phosphorylation) as has been previous suggested (Coste et al., 2015). There is evidence that cytoskeletal interaction is important in modulating mechanogating of Piezo1, with removal of the cytoskeleton leading to a decrease in the pressure required for channel opening (Cox et al., 2016).

Our simulations suggest that membrane cholesterol composition regulates the dimensions of the dome created by Piezo1, with dome depth converging on a minimum at 10% cholesterol in the truncated version of Piezo1 and at 10% and 20% in the full-length model of Piezo1 and increasing at smaller or greater cholesterol concentrations. Our experimental results are in good agreement with this relationship, with Piezo1 channel activity at a peak in the native cholesterol composition of HUVECs and diminishing as cholesterol is added or removed. At this stage it is unclear how the native cholesterol concentration of the HUVEC plasma membrane relates to the cholesterol concentrations used in our simulations. Published data suggests that cholesterol comprises 21% of total HUVEC lipids (Murphy et al., 1992). Our analysis suggests that some of the interactions of Piezo1 with cholesterol occur in CRAC and CARC motifs. It should be noted, however, that less than half of the CRAC or CARC on Piezo1 had significant cholesterol interactions. This may be due to the fact that although the interaction of CRAC and CARC motifs with cholesterol has been shown in a number of systems (Fantini and Barrantes, 2013), their predictive value for cholesterol binding is limited due to the loose definition of the consensus sequence (Palmer, 2004). Experimentally verifying the predictive value of these motifs would require mutating multiple stretches of amino acids in association with functional studies involving cholesterol. Such extensive changes could potentially have substantial structural effects, which would make it challenging to interpret results.

The ability of proteins to modify their local lipid environment has been described both computationally (Corradi et al., 2018) and experimentally (Prabudiansyah et al., 2015). Uniquely, in addition to its effect on local membrane curvature, Piezo1 changes its local lipid environment, by forming an annulus of PIP_2_ through strong preferential interaction. This is a demonstration of how a curved ion channel changes its local membrane environment both in terms of its lipid composition and of its topology. The PIP_2_ annulus around Piezo1 could affect PIP_2_ signaling by creating a hotspot around Piezo1 or acting as a PIP_2_ sink to prevent PIP_2_ signaling further way. This is especially so for the Piezo1_full_ model, where there is a high number of PIP_2_ contacts with the modelled W1806 – A1951 loop and N-terminal residues. Moreover, the annulus formed around Piezo1_full_ has the additional feature of crossing the interface between blades of adjacent chains, as well as relatively depleting PIP_2_ around the Piezo1 core. The concentration of PIP_2_ at the interface between Piezo1 blades may serve to stabilize their position, and relative paucity of PIP_2_ around the core may tune PIP_2_ sensitivity in this region to lower PIP_2_ concentrations. One site which could be thus regulated is K2183. This site is homologous to K2167 on human Piezo1, which is the site of an in-frame deletion, hPiezo1 K2166-2169del, which causes dehydrated hereditary stomatocytosis (DHS) (Andolfo et al., 2013). PIP_2_ has been reported to regulate Piezo1 channel activity experimentally (Borbiro et al., 2015). Our data raises the possibility that this mutation may cause DHS through loss of PIP_2_ regulation.

In summary, this study proposes a 3D structure for the full-length Piezo1 channel, which has proven challenging to obtain experimentally. The full-length structure is critical when studying Piezo1 as it modifies the unique footprint of Piezo1 in a lipid bilayer. Piezo1 unique footprint has a unique trilobed topology with specific interactions with lipids e.g. PIP_2_. The footprint and Piezo1 function are modified by cholesterol, which could be important in atherosclerotic disease, a process in which both cholesterol and endothelial shear stress sensing, which Piezo1 contributes to, play a key role. Therefore, our findings suggest novel ways by which Piezo1 could act as an integrator of mechanical responses in health and disease.

## MATERIALS AND METHODS

### Molecular modelling of Piezo1_trunc_

Missing loops were modelled on the published mPiezo1 structure (PDB: 6B3R) (Guo and MacKinnon, 2017) using Modeller (v9.19) (Šali and Blundell, 1993). Five models were generated, and the best model selected using the discrete optimized protein energy (DOPE) method (Shen and Sali, 2006). Loops which remained unstructured after modelling were removed from the model before simulation. These loops were located at residues 718 – 781, 1366 – 1492, 1579 – 1654, and 1808 – 1951.

### Molecular modelling of Piezo1_full_

Structural data were obtained from the published cryo-EM structure (PDB: 6B3R). Missing residues were added with MODELLER (v 9.19) and the loop refinement tool (Fiser et al., 2000) was used to remove a knot in one chain between residues 1490-1498. The best model was selected out of 10 candidates according to the DOPE method. In order to model the missing 3 N-terminal bundles from the template 6B3R (i.e. residues 1-576), we first carried out a transmembrane and structural prediction using MEMSAT-SVM (Nugent and Jones, 2012) and PSIPRED webserver (Buchan and Jones, 2019; Jones, 1999). As a structural template we used the bundles 4-5-6 from a Piezo1 blade (i.e. residues 577-1129) in combination with MODELLER. The PSI/TM-Coffee web tool in slow/accurate homology extension mode was used to obtain the target-template alignment. The final target-template alignment was carefully checked and manually modified to avoid fragmentation of secondary structure elements and transmembrane helices. During modeling we imposed canonical α-helix conformations for the residues 2-12, 97-103 and 183-189. The loop modeling routine of MODELLER (Fiser et al., 2000) was used to remove a knot between residues 149-182 and 294-317 selecting the best loop out of 5 according the DOPE score. The position of the obtained bundles 1-2-3 with respect to the rest of the protein was obtained by superposing the bundle 3 to the bundle 4 from a single chain using UCSF Chimera (Pettersen et al., 2004). Subsequently, bundles 1-2-3 were manually moved to avoid superposition. This procedure ensured that the new modeled bundles 1-2-3 followed a similar direction of the partial Piezo1 blade resolved by cryo-EM. We then used UCSF Chimera to impose canonical α-helix conformation to cytoplasmic residues 567-587, 747-752, 776-806, 1420-1424, 1437-1446, 1449-1468, 1632-1635, 1645-1650 and 1926-1968 as predicted by PSIPRED. This full-length Piezo1 chain was superposed to the others in 6B3R using UCSF Chimera obtaining the trimeric Piezo1 structure. The Piezo1 full-length model obtained was energy minimized in vacuum with GROMACS 5.0.7 (Abraham et al., 2015) prior simulations.

### Coarse-grained simulations

The Piezo1 models obtained were converted to a coarse-grained resolution using the *martinize* script (De Jong et al., 2012) and further energy minimized in vacuum with GROMACS 5.0.7. The CG-MD simulations were performed using the Martini 2.2 force field (De Jong et al., 2012) and GROMACS 5.0.7. To model the protein secondary and tertiary structure an elastic network model with a cut-off distance of 7 Å was used. The elastic network restricts any major conformational change within the protein during the CG-MD simulations. For the equilibration simulation Piezo1 models were inserted in a complex asymmetric bilayer using the INSert membrANE tool (Wassenaar et al., 2015). The system compositions are described in Supplementary Table 1. All systems were initially assembled in a simulation box of size 44 × 44 × 24 nm. The systems were neutralized with a 150 mM concentration of NaCl. The models were further energy minimized and subsequently equilibrated with the protein particles restrained (1000 kJmol_-1_nm_-2_) to allow the membrane bilayer to equilibrate around the model. Equilibration time was 500 ns for Piezo1_full_, 100 ns for Piezo1_trunc_. These long CG-MD equilibration steps were essential to equilibrate the lipid bilayer around Piezo1 and reconstitute the dome previously hypothesized (Guo and MacKinnon, 2017). All simulations were performed at 323 K, with protein, lipids and solvent separately coupled to an external bath using the V-rescale thermostat (Bussi et al., 2007) (coupling constant of 1.0). Pressure was maintained at 1 bar (coupling constant of 1.0) with semi-isotropic conditions and compressibility of 3 × 10^-6^ using the Berendsen barostat (Berendsen et al., 1984) during equilibrations and the Parrinello-Rahman barostat (Parrinello and Rahman, 1981) during productions. Lennard-Jones and Coulombic interactions were shifted to zero between 9 and 12 Å, and between 0 and 12 Å, respectively. After the equilibration phase of the Piezo1 full-length, we removed lipid molecules that flipped between leaflets in order to restore the membrane asymmetry. After this step, the systems were further energy minimized and when applicable, neutralized with counterions. A preliminary run of 5 ns using an integration step of 10 fs was carried out prior the production phase. For each Piezo1 model five unrestrained repeat simulations of 3 μs each were run using an integration step of 20 fs.

### Piezo1 footprint depth analysis

A Python script was written to measure the depth of the Piezo1 dome. First, the simulation trajectory is fitted to the protein coordinates using the GROMACS tool gmx trjconv. The coordinates of CG phosphate head group residues in each frame of the fitted trajectory are extracted by a Python script which performs the following analysis. The phosphate atoms are then used to separate the bilayer leaflets bilayers using a branching network algorithm. Briefly, this method starts with a single particle, and identifies the other head group particles which are within a cutoff distance (2 nm) of the starting residue. The cutoff distance is selected to be smaller than the separation between the bilayer leaflets. Head group particles identified in this way are added to the same leaflet as the starting residue. This process iterates repeatedly until no more new particles can be added, and remaining residues are assumed to be part of the other leaflet. The process is then repeated starting in the other leaflet to confirm that the leaflet identification is correct.

For each leaflet, the depth of the dome is calculated as the difference between the surface level and the bottom of the dome. The surface level is taken to be the average z-coordinate for the head group residues with z-coordinate in excess of the 90_th_ centile. The average is used here to minimise the effect of random fluctuation of the membrane. For the bottom of the dome, the z-coordinate of the head group residue with the absolute lowest z-coordinate is used. This is because the bottom of the dome is prone to far less fluctuation, being fixed to Piezo1, which in turn is the fitting reference for the rest of the simulation.

To generate the height map of the bilayer leaflets, for each frame, CG phosphate beads were binned along the x and y axes; 75 bins for each axis. The average z-coordinate of beads contained in each bin was calculated and stored in a matrix for each frame. The matrices of all frames are averaged to create the final height map, which is plotted using the PyPlot library.

Code used is available at: https://github.com/jiehanchong/membrane-depth-analysis

### Analysis of protein and lipid density

The repeat simulations for each system were concatenated and fitted to the Piezo1 pore (residues 2105-2547). gmx densmap from the GROMACS package was used to generate 2-dimensional density map for protein and individual lipid species, with summation of density along the z-axis.

### Analysis of protein-lipid contacts

Alll 5 repeat simulations were concatenated for each system. Contact between CG lipid head group beads and protein beads were calculated using gmx mindist from the GROMACS package. The results were checked to ensure consistent results between each chain in the trimer. The contacts for individual Piezo1 chains were added together. Data was plotted using Grace (http://plasma-gate.weiz-mann.ac.il/Grace/).

### Cell culture

HUVECs were purchased from Lonza and cultured in Endothelial Cell Basal Medium (EBM-2) supplemented with 2% fetal calf serum (Sigma) and the following growth factors: 10 ng.ml_- 1_ vascular endothelial growth factor, 5 ng.ml^-1^ human basic fibroblast growth factor, 1 μg.ml^-1^ hydrocortisone, 50 ng.ml^-1^ gentamicin, 50 ng.ml^-1^ amphotericin B, and 10 μg.ml^-1^ heparin. These growth factors were supplied as a bullet kit (Cell Media and Bullet Kit, Lonza). HUVECs were passaged 2-6 times.

HEK 293 cells stably expressing human Piezo1 (P1) under a tetracycline inducible promoter or stably expressing mouse Piezo1 without need for tetracycline induction were utilised. Cells used for Yoda1 experiments were cultured in Dulbecco’s modified Eagle’s medium-F12 GlutaMAX (Invitrogen, Paisley, UK). Cells used in patch clamp experiments were cultured in Dulbecco’s Modified Eagle’s medium (Invitrogen). All HEK cell culture media were supplemented with 10% foetal calf serum (Sigma) and 1% Pen/Strep (Sigma-Aldrich). For patch clamp experiments, the cells were plated on poly-lysine-coated coverslips in bath solution (see Patch-clamp recording) 1 hr before experiments.

All cells were maintained at 37 °C in a humidified atmosphere containing 5 % CO_2_.

### Short interfering (si) RNAs

HUVECs were transfected at 90 % confluence with 50 nM siRNA using Lipofectamine 2000 in OptiMEM (Gibco) as per the manufacturer’s instructions (Invitrogen). Medium was replaced after 3-4 hr and cells were used for experimentation 48 hr post transfection. siRNA was provided from Ambion. The sequence for Piezo1 is GCCUCGUGGUCUACAAGAUtt. Scrambled negative control siRNA provided by Ambion was used as a control.

### Intracellular Ca_2+_ measurement

Cells were incubated with fura-2AM (2 μM) (Thermo Fisher Scientific) for 60 min at 37 °C followed by a 30 min wash at room temperature. All cholesterol treatments were added prior to the 30 min wash, for 30 min at 37 °C, and were present during Ca_2+_-measurements. Measurements were made at room temperature on a 96-well plate reader (FlexStation, Molecular Devices) or a Zeiss Axiovert fluorescent microscope equipped with x20 (NA 0.75) Fluar objective. The change (Δ) in intracellular Ca_2+_ concentration was indicated as the ratio of fura-2 emission (510 nm) intensities for 340 nm and 380 nm excitation (F340/380). The recording solution (standard bath solution, SBS) contained (mM): 130 NaCl, 5 KCl, 8 D-glucose, 10 HEPES, 1.2 MgCl_2_, 1.5 CaCl_2_, titrated to pH 7.4 with NaOH.

### Cholesterol treatment

Water soluble cholesterol in poly(ethylene glycol) [PEG]-600, cholesterol in complex with methyl β cyclodextrin (cholesterol at 40mg/g), methyl β cyclodextrin were purchased from Sigma. Cells were incubated with cholesterol enriching or depleting agents for 30 minutes, 37°C, 0% CO_2_ in SBS.

### Chemical reagents

Yoda1 (Tocris) was prepared in stocks at 10 mM in DMSO (Sigma).

### Patch-clamp recording

Macroscopic membrane currents through outside-out patches were recorded using standard patch-clamp technique in voltage-clamp mode at room temperature (21-23 °C). The holding voltage was −80 mV. Patch pipettes were fire-polished and had resistance of 4–7MΩ when filled with pipette solution. The bath and pipette solutions were identical and contained (in mM): NaCl 140, HEPES 10 and EGTA 5 (pH 7.4, NaOH). Recordings were made using an Axopatch-200B amplifier (Axon Instruments, Inc., USA) equipped with Digidata 1550B and pClamp 10.6 software (Molecular Devices, USA). Pressure steps of 200-ms duration were applied to the patch pipette using a High Speed Pressure Clamp HSPC-1 System (ALA Scientific Instruments, USA). Current records were analogue filtered at 1 kHz and digitally sampled at 5 kHz.

### Statistical analysis

The study was aimed at discovering components of a biological mechanism using various cell/molecular studies to address a single hypothesis. In the absence of prior knowledge of the mechanism, power calculations were not considered to be applicable. We selected numbers of independent repeats of experiments base on prior experience of studies of this type. In all cases the number of independent repeats was at least 4.

For Yoda1 experiments, OriginPro 9.1 was used for data analysis and graph production. Averaged data are presented as mean±s.e.mean.

For patch clamp experiments, data were analysed and plotted using pClamp 10.6 and MicroCal Origin 2018 (OriginLab Corporation, USA). Averaged data are presented as mean±s.d.mean.

For both sets of experiments, most data were produced in pairs (test and control) and these data pairs were compared statistically using *t* tests. One-way ANOVA followed by Tukey posthoc test was used for comparing multiple groups. Statistically significant difference is indicated by * (*P*<0.05) and no significant difference by NS (*P*>0.05). The number of independent experiments is indicated by n.

Outlying data were not detected or excluded. The number of replicates per independent experiment was 4 for Ca_2+_ assays and 1 for patch-clamp assays.

## ACKNOWLEDGEMENTS

JC is funded by an MRC Clinical Research Training Fellowship, DJB by a Wellcome Trust Investigator Award, AJH by a BBSRC PhD Studentship, JS by a BHF Intermediate Basic Science Research Fellowship, and ACK and DDV by an Academy of Medical Sciences and Wellcome Trust Springboard Award.

## Supplementary information

**Supplementary Table 1:** Membrane lipid compositions used for the CG-MD simulations

| Name | Extracellular leaflet |  |  |  | Cytoplasmic leaflet |  |  |  |  |
| --- | --- | --- | --- | --- | --- | --- | --- | --- | --- |
|  | PC | PE | SM | CHOL | PC | PE | PS | PIP2 | CHOL |
| CHOL0<sub>trunc</sub> | 75 | 20 | 5 | 0 | 70 | 20 | 5 | 5 | 0 |
| CHOL5<sub>trunc</sub> | 70 | 20 | 5 | 5 | 65 | 20 | 5 | 5 | 5 |
| CHOL10<sub>trunc</sub> | 65 | 20 | 5 | 10 | 60 | 20 | 5 | 5 | 10 |
| CHOL20<sub>trunc</sub> | 55 | 20 | 5 | 20 | 50 | 20 | 5 | 5 | 20 |
| CHOL30<sub>trunc</sub> | 45 | 20 | 5 | 30 | 40 | 20 | 5 | 5 | 30 |
| CHOL40<sub>trunc</sub> | 35 | 20 | 5 | 40 | 30 | 20 | 5 | 5 | 40 |
| CHOL0<sub>full</sub> | 75 | 20 | 5 | 0 | 70 | 20 | 5 | 5 | 0 |
| CHOL10<sub>full</sub> | 65 | 20 | 5 | 10 | 60 | 20 | 5 | 5 | 10 |
| CHOL20<sub>full</sub> | 55 | 20 | 5 | 20 | 50 | 20 | 5 | 5 | 20 |
| CHOL30<sub>full</sub> | 45 | 20 | 5 | 30 | 40 | 20 | 5 | 5 | 30 |

Lipid composition is expressed in molar percentages

**Supplementary Figure 1:**
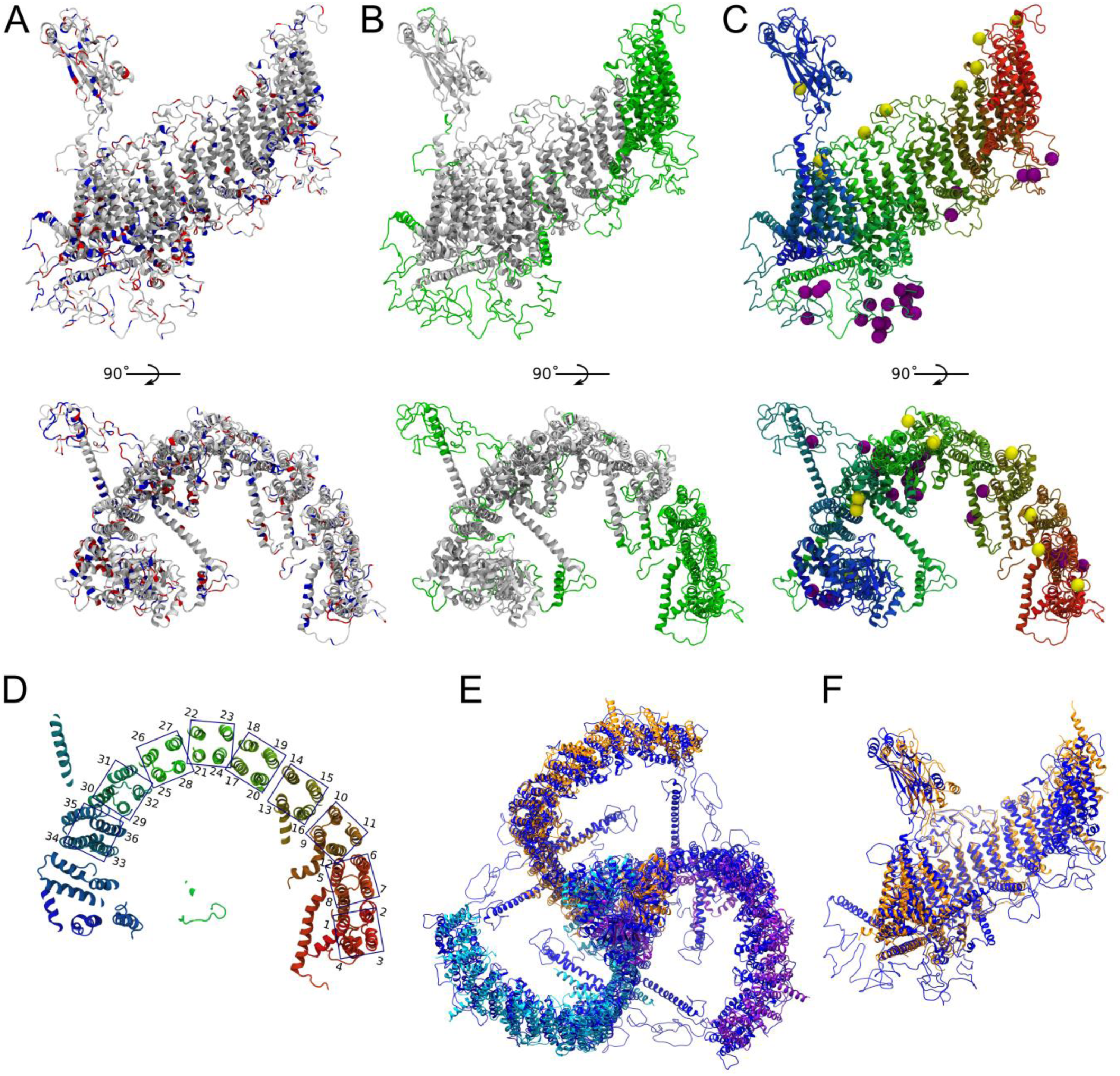
Characterization of a single full-length chain Piezo1 mouse model. **(A)** A single full-length chain is shown as ribbon with positively (His, Lys, Arg) and negatively (Asp, Glu) charged residues colored in blue and red, respectively. **(B)** A single full-length chain is shown as ribbon with modelled missing residues from the PDB 6 B3R, highlighted in green. **(C)** A single full-length chain is shown as ribbon and colored rainbow from N-terminal (red) to C-terminal (blue). Spheres are Cα atoms from residues previously identified as exposed to the extracellular side (Myc tagged, yellow) and phosphorylated (purple) (Coste et al., 2015). Myc tagged residues are: 102, 304, 508, 669, 897, 1071, 1765, 2071, 2075 and 2336. Phosphorylated residues are: 351, 396, 397, 399, 738, 758, 1385, 1389, 1390, 1593, 1600, 1604, 1608, 1610, 1612, 1617, 1626, 1627, 1631, 1640, 1644, 1646 and 1837. **(D)** Cropped visualization of a single Piezo1 full-length chain. The numbering and position of each of the 36 bundles is indicated. **(E)** Structural superposition between the Piezo1 full-length model (blue) and the Piezo2 cryo-EM structure (chains are in orange, purple and cyan; PDB: 6KG7) performed with UCSF Chimera over the Cα atoms. **(F)** A single chain from the Piezo2 cryo-EM structure (orange) and a Piezo1 full-length model superposed with UCSF Chimera over their Cα atoms. The long loops in the cytoplasm (residues 718-781, 1366-1492, 1579-1654, 1808-1951, 542-568, 337-418, 145-182) were not considered in this alignment.

**Supplementary Figure 2:**
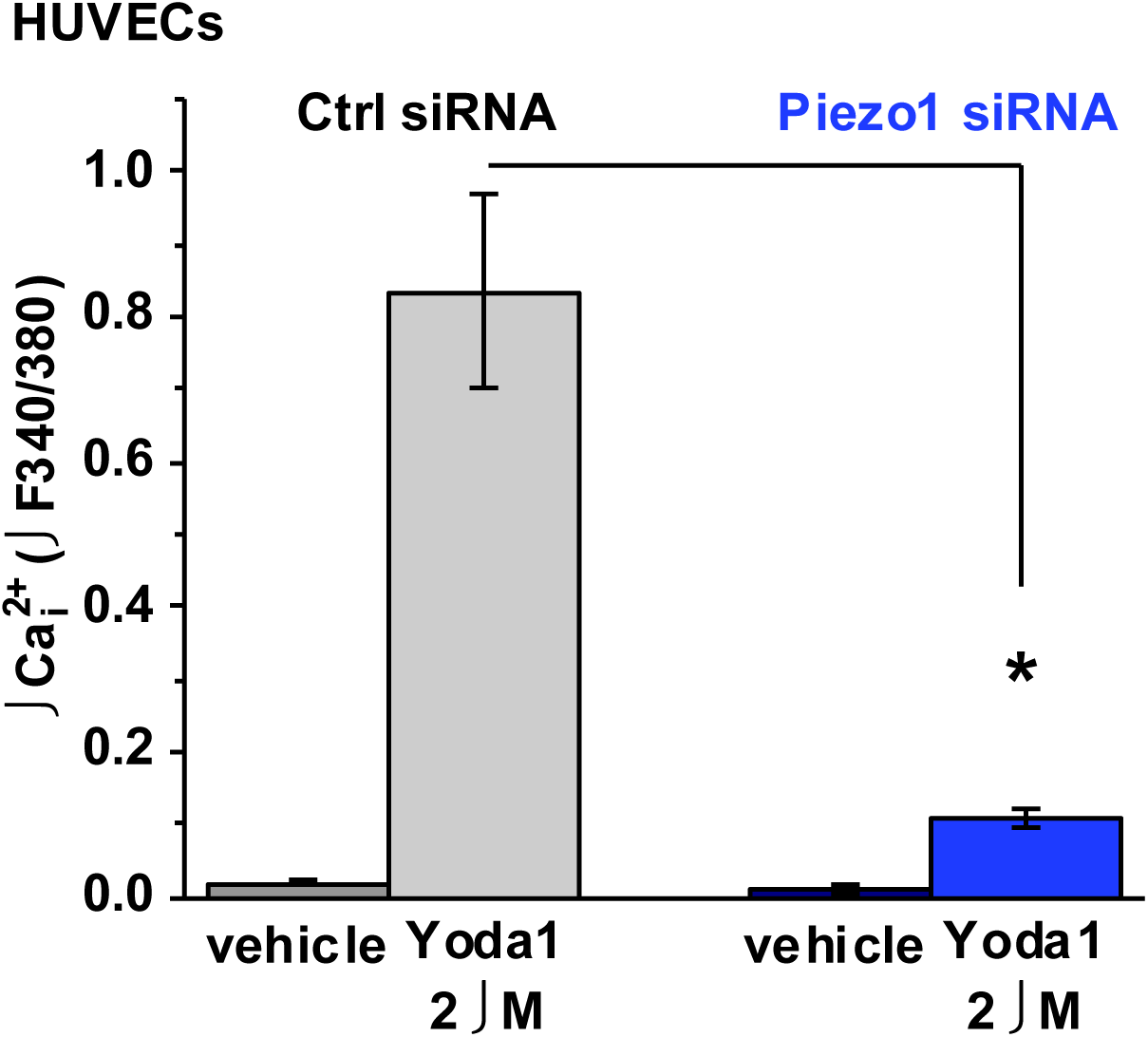
Yoda1 specifically activates Piezo1 in human umbilical vein endothelial cells (HUVECs). Shown are mean ± s.e.mean data for the change (Δ) in the intracellular Ca^2+^ concentration evoked by vehicle (dimethylsulphoxide) or vehicle plus 2 μM Yoda1 in HUVECs transfected with either control siRNA (Ctrl siRNA) or Piezo1 siRNA (n=3 for each). The Yoda1 response was significantly smaller in the Piezo1 siRNA group (P<0.05).

**Supplementary Figure 3:**
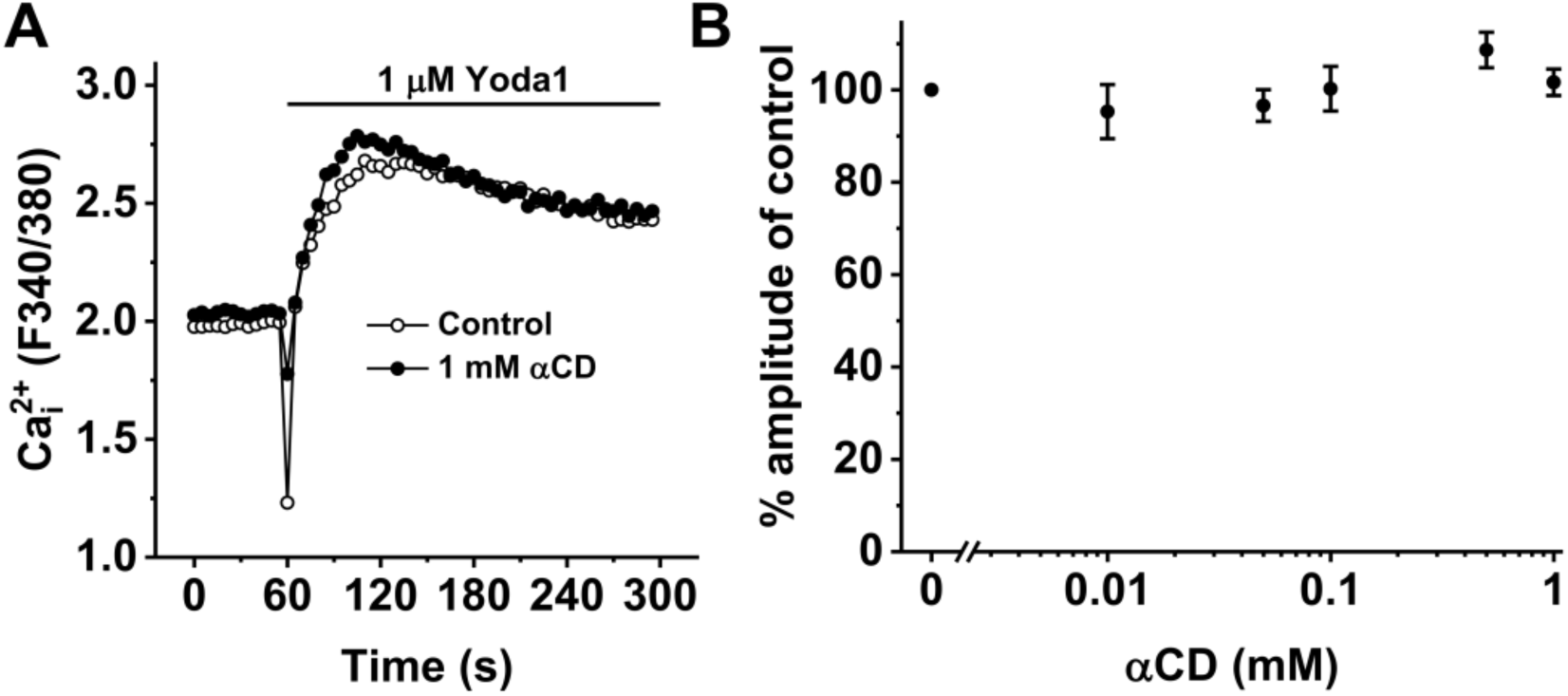
Alpha cyclodextrin (αCD) has no effect on Piezo1 activation. **(A)** an example measurement of the change in intracellular Ca^2+^ concentration evoked by 1 μM Yoda1 in HUVECs pre-treated with 0 (control) and 1 mM αCD. **(B)** mean data of peak amplitudes of HUVEC responses to increasing doses of αCD (1-0.01 mM). n/N=3/4.

**Supplementary Table 2:** Piezo1_trunc_ residues with cholesterol contacts exceeding cutoff of 60% of maximum contacts.

|  |  |  |  |
| --- | --- | --- | --- |
| <u>Y582</u><sub>e</sub> | <u>Q703</u><sub>e</sub> | D1211<sub>cef</sub> | <u>M1731</u><sub>cef</sub> |
| <u>V583</u><sub>def</sub> | <u>H705</u><sub>f</sub> | L1215<sub>ef</sub> | <u>T1732</u><sub>ce</sub> |
| V589<sub>def</sub> | <u>Y706</u><sub>ef</sub> | M1494<sub>f</sub> | <u>V1735</u><sub>c</sub> |
| <u>T619</u><sub>e</sub> | <u>Y782</u><sub>f</sub> | M1495<sub>f</sub> | E1738<sub>f</sub> |
| <u>Q622</u><sub>bcdef</sub> | <u>L786</u><sub>def</sub> | A1499<sub>f</sub> | <u>M1740</u><sub>bcde</sub> |
| <u>T626</u><sub>ef</sub> | <u>S789</u><sub>def</sub> | Q1503<sub>ef</sub> | <u>T1743</u><sub>bdf</sub> |
| <u>L627</u><sub>bcde</sub> | <u>F800</u><sub>ef</sub> | **F1504**<sub>bcdef</sub> | <u>K1744</u><sub>bcd</sub> |
| <u>W636</u><sub>ef</sub> | H808<sub>cf</sub> | <u>L1508</u><sub>ef</sub> | <u>L1774</u><sub>d</sub> |
| <u>W637</u><sub>def</sub> | <u>K811</u><sub>cdef</sub> | G1509<sub>ce</sub> | G1775<sub>b</sub> |
| L647<sub>e</sub> | <u>F986</u><sub>e</sub> | <u>T1512</u><sub>e</sub> | <u>K1784</u><sub>ef</sub> |
| V650<sub>e</sub> | **A1019**<sub>bcdef</sub> | R1671<sub>cdef</sub> | <u>Y1785</u><sub>def</sub> |
| Y651<sub>ef</sub> | **T1022**<sub>bcdef</sub> | T1672<sub>cef</sub> | F2028<sub>e</sub> |
| F655<sub>ef</sub> | **R1023**<sub>bcdef</sub> | Y1680<sub>cef</sub> | <u>W2060</u><sub>ef</sub> |
| F687<sub>f</sub> | <u>Y1062</u><sub>ef</sub> | A1706<sub>d</sub> | V2141<sub>f</sub> |
| A699<sub>e</sub> | <u>Y1190</u><sub>f</sub> | <u>W1717</u><sub>f</sub> | W2142<sub>ef</sub> |
| <u>L702</u><sub>def</sub> | V1208<sub>ef</sub> | <u>R1728</u><sub>cef</sub> | <u>Y2189</u><sub>cdef</sub> |

The simulated cholesterol concentration in which the residue exceeds cutoff is represented by superscript letters after each residue. Key: b – 5%, c – 10%, d – 20%, e – 30%, f – 40%. Residues which were above cut-off at all simulated cholesterol concentrations are in bold. Residues coinciding with CRAC or CARC motifs are shown in blue. Residues also exceeding cut-off in Piezo1<sub>full</sub> simulations are underlined.

**Supplementary Table 3:** Piezo1_full_ residues with cholesterol contacts exceeding cut-off of 60% of maximum contacts.

|  |  |  |  |  |  |
| --- | --- | --- | --- | --- | --- |
| **L20**<sub>cde</sub> | H74<sub>e</sub> | **T209**<sub>cde</sub> | L778<sub>de</sub> | T1512<sub>e</sub> | F1932<sub>e</sub> |
| **R26**<sub>cde</sub> | A76<sub>de</sub> | V233<sub>e</sub> | **R782**<sub>cde</sub> | V1513<sub>e</sub> | <u>Q1944</u><sub>e</sub> |
| L32<sub>de</sub> | F77<sub>e</sub> | L235<sub>e</sub> | L786<sub>e</sub> | **R1671**<sub>cde</sub> | I1988<sub>e</sub> |
| V33<sub>e</sub> | C80<sub>e</sub> | A236<sub>e</sub> | S789<sub>e</sub> | A1706<sub>de</sub> | F2028<sub>e</sub> |
| P42<sub>ce</sub> | I105<sub>e</sub> | C261<sub>e</sub> | <u>F800</u><sub>e</sub> | L1716<sub>e</sub> | <u>W2060</u><sub>e</sub> |
| W43<sub>ce</sub> | **A123**<sub>cde</sub> | G265<sub>e</sub> | A1019<sub>ce</sub> | W1717<sub>ce</sub> | <u>F2130</u><sub>e</sub> |
| **P47**<sub>cde</sub> | **P124**<sub>cde</sub> | **T313**<sub>cde</sub> | **T1022**<sub>cde</sub> | M1740<sub>ce</sub> | <u>Y2189</u><sub>e</sub> |
| R58<sub>de</sub> | R187<sub>e</sub> | W533<sub>e</sub> | **R1023**<sub>cde</sub> | T1743<sub>e</sub> |  |
| **R61**<sub>cde</sub> | R188<sub>e</sub> | <u>Q622</u><sub>de</sub> | Y1190<sub>e</sub> | K1744<sub>e</sub> |  |
| A62<sub>e</sub> | T205<sub>e</sub> | <u>L627</u><sub>e</sub> | L1508<sub>e</sub> | <u>K1784</u><sub>e</sub> |  |
| C65<sub>de</sub> | S206<sub>e</sub> | <u>W637</u><sub>e</sub> | G1509<sub>e</sub> | <u>Y1785</u><sub>e</sub> |  |

The simulated cholesterol concentration in which the residue exceeds cut-off is represented by superscript letters after each residue. Key: c – 10%, d – 20% and e – 30%. Residues which were above cut-off at all simulated cholesterol concentrations are in bold. Residues coinciding with CRAC or CARC motifs are shown in blue. Residues also exceeding cut-off in Piezo1<sub>trunc</sub> simulations are underlined.

**Supplementary Table 4:**
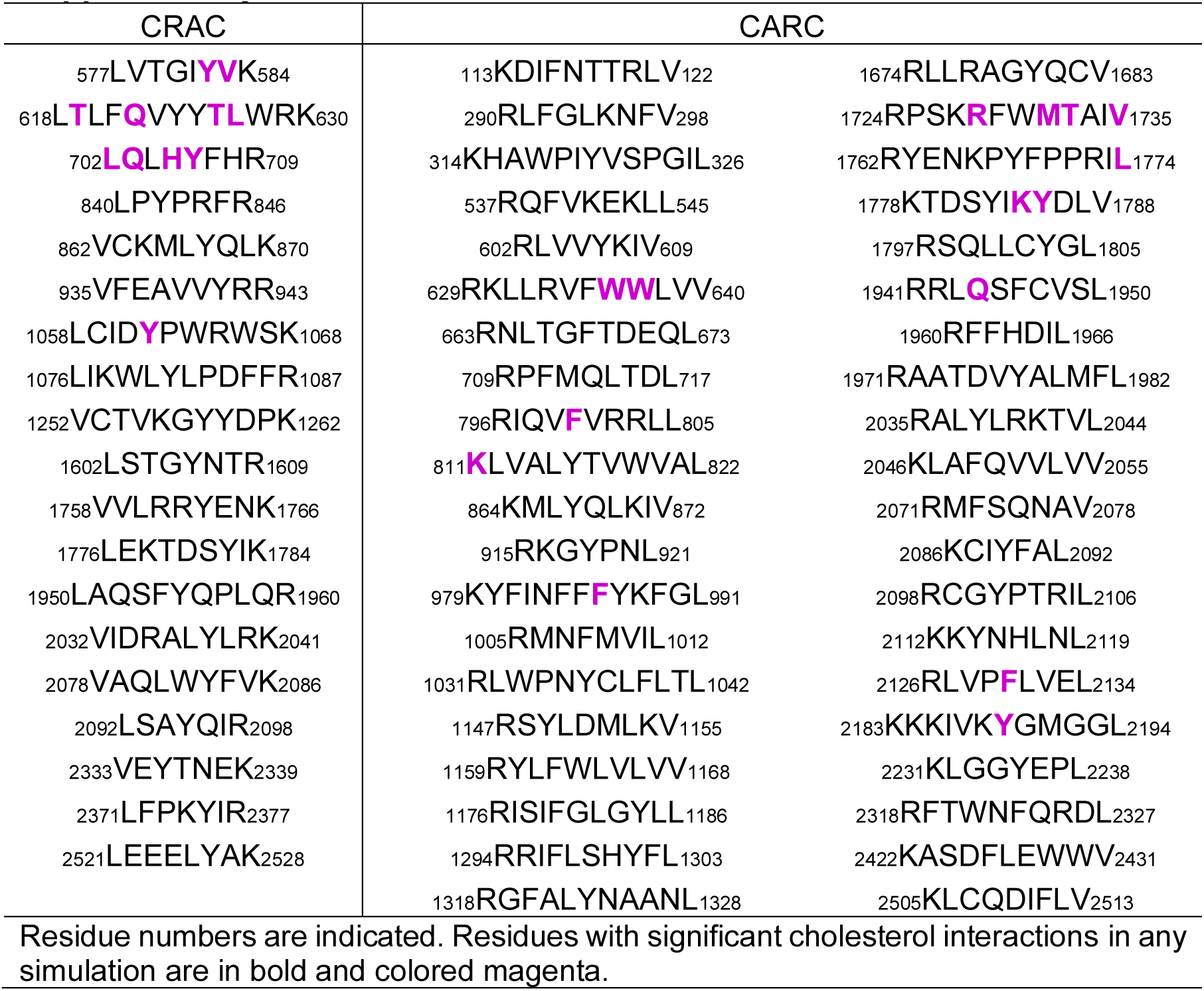
CRAC and CARC motifs in mPiezo1

**Supplementary Table 5:** Piezo1_trunc_ residues with PIP_2_ contacts exceeding cut-off of 70% of maximum contacts.

|  |  |  |
| --- | --- | --- |
| L577<sup>ef</sup> | **<u>R844</u>**<sub>abcdef</sub> | Q1496<sup>e</sup> |
| T579<sub>acdef</sub> | **<u>R846</u>**<sub>abcdef</sub> | **R1497**<sub>abcdef</sub> |
| **K584**<sub>abcdef</sub> | **R942**<sub>abcdef</sub> | K1727<sub>abcde</sub> |
| L627<sub>cef</sub> | R949<sub>bcd</sub>e | R1728<sub>abce</sub> |
| **W628**<sub>abcdef</sub> | **R1023**<sub>abcdef</sub> | Q1952<sup>b</sup> |
| **R629**<sub>abcdef</sub> | **R1024**<sub>abcdef</sub> | R2040<sub>acd</sub> |
| **K630**<sub>abcdef</sub> | **R1025**<sub>abcdef</sub> | K2183<sub>acde</sub> |
| **R633**<sub>abcdef</sub> | **R1031**<sub>abcdef</sub> | K2184<sub>abde</sub> |
| H708<sup>df</sup> | K1201<sup>c</sup> | K2185<sub>acde</sub> |
| **R709**<sub>abcdef</sub> | **R1204**<sub>abcdef</sub> | G2188<sup>c</sup> |
| **R782**<sub>abcdef</sub> | **H1493**<sub>abcdef</sub> | R2517<sub>cd</sub> |

The simulated cholesterol concentration of systems in which the residue exceeds the cutoff is indicated by superscript letters as follow: a – 0%, b – 5%, c – 10%, d – 20%, e – 30%, f – 40%. Residues which were above the cutoff in all the Piezo1<sub>trunc</sub> systems are in bold. Residues also exceeding cut-off in Piezo1<sub>full</sub> simulations are underlined.

**Supplementary Table 6:** Piezo1full residues with PIP2 contacts exceeding cut-off of 70% of maximum contacts.

|  |  |  |  |
| --- | --- | --- | --- |
| T56<sub>ce</sub> | **R466**<sub>acde</sub> | **R1024**<sub>acde</sub> | R1924<sub>acd</sub> |
| **G57**<sub>acde</sub> | **R629**<sub>acde</sub> | **R1025**<sub>acde</sub> | H1925<sub>ac</sub> |
| **R58**<sub>acde</sub> | **K630**<sub>acde</sub> | <u>K1201</u><sub>ad</sub> | **K1929**<sub>acde</sub> |
| L60<sub>ac</sub> | **L632**<sub>acde</sub> | **R1204**<sub>acde</sub> | S1930<sub>cd</sub> |
| **R61**<sub>acde</sub> | **R633**<sub>acde</sub> | R1891<sub>acd</sub> | K1931<sub>acd</sub> |
| **R144**<sub>acde</sub> | F711<sub>cde</sub> | R1892<sub>acd</sub> | **R1933**<sub>acde</sub> |
| R151<sub>e</sub> | G724<sub>ce</sub> | R1893<sub>acd</sub> | **R1935**<sub>acde</sub> |
| **K186**<sub>acde</sub> | **R726**<sub>acde</sub> | K1894<sub>cd</sub> | **K1937**<sub>acde</sub> |
| **R187**<sub>acde</sub> | **R844**<sub>acde</sub> | **R1920**<sub>acde</sub> | R1941<sub>acd</sub> |
| R348<sub>de</sub> | <u>R846</u><sub>c</sub> | **K1921**<sub>acde</sub> | R1942<sub>acd</sub> |
| K349<sub>e</sub> | **R1023**<sub>acde</sub> | R1922<sub>cd</sub> |  |

The cholesterol concentration of systems in which the residue exceeds the cut-off is indicated by superscript letters as follow: a – 0%, c – 10%, d – 20% and e – 30%. Residues which were above the cut-off in all the Piezo1<sub>full</sub> systems are in bold. Residues also exceeding cut-off in Piezo1<sub>trunc</sub> simulations are underlined.

**Supplementary Figure 4:**
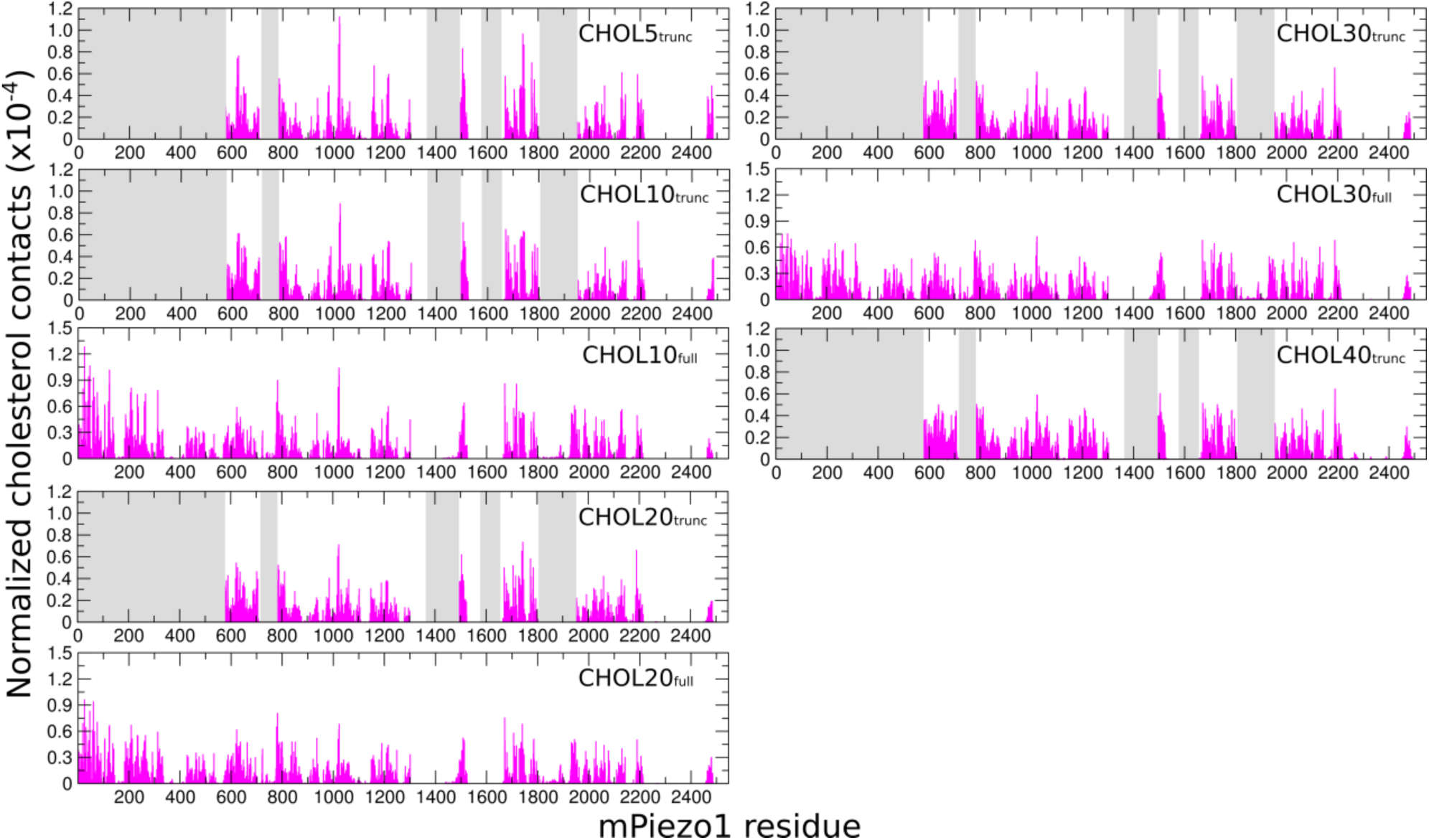
Histograms of Piezo1-cholesterol contacts in all simulations. Each histogram is labelled with the corresponding simulation. Grey bars represent the residues missing from Piezo1_trunc_.

**Supplementary Figure 5:**
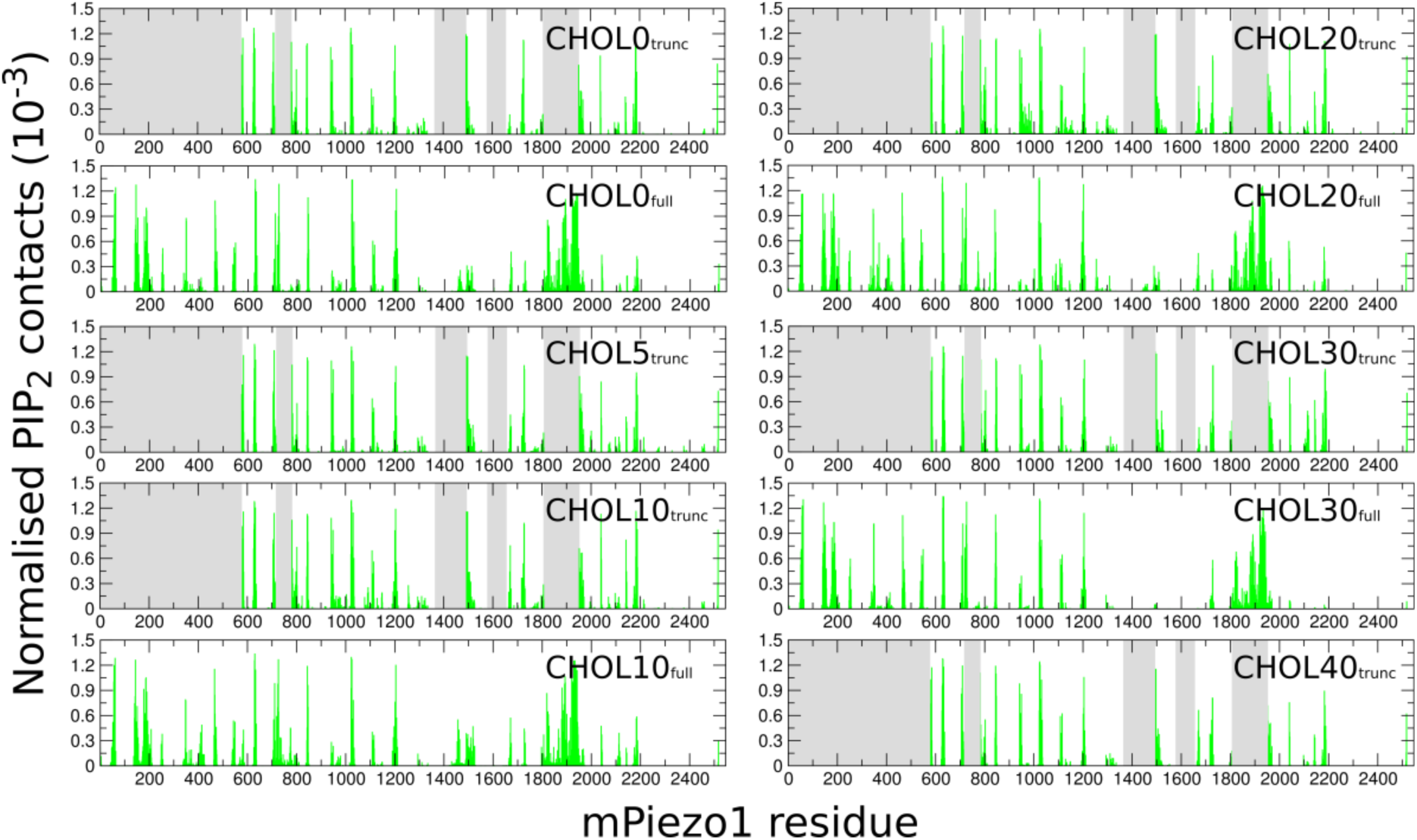
Histograms of Piezo1-PIP_2_ contacts in all simulations. Each histogram is labelled with the corresponding simulation. Grey bars represent the residues missing from Piezo1_trunc_.

**Supplementary Figure 6:**
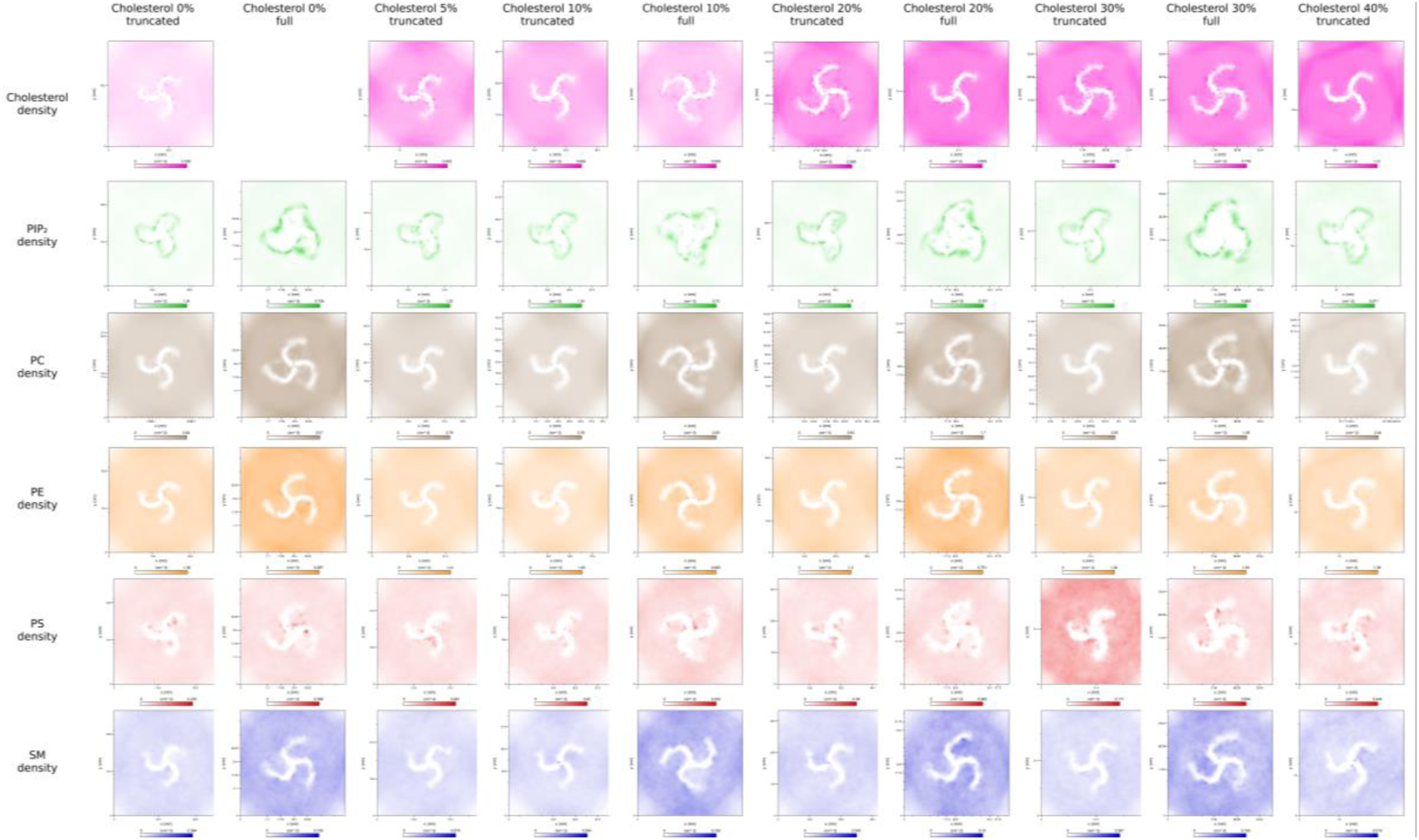
Matrix of lipid density maps for all simulations conducted. Lipids are color coded as cholesterol – magenta, PIP_2_ – green, POPC – tan, POPE – orange, POPS – red, SM – blue.

**Supplementary Figure 7:**
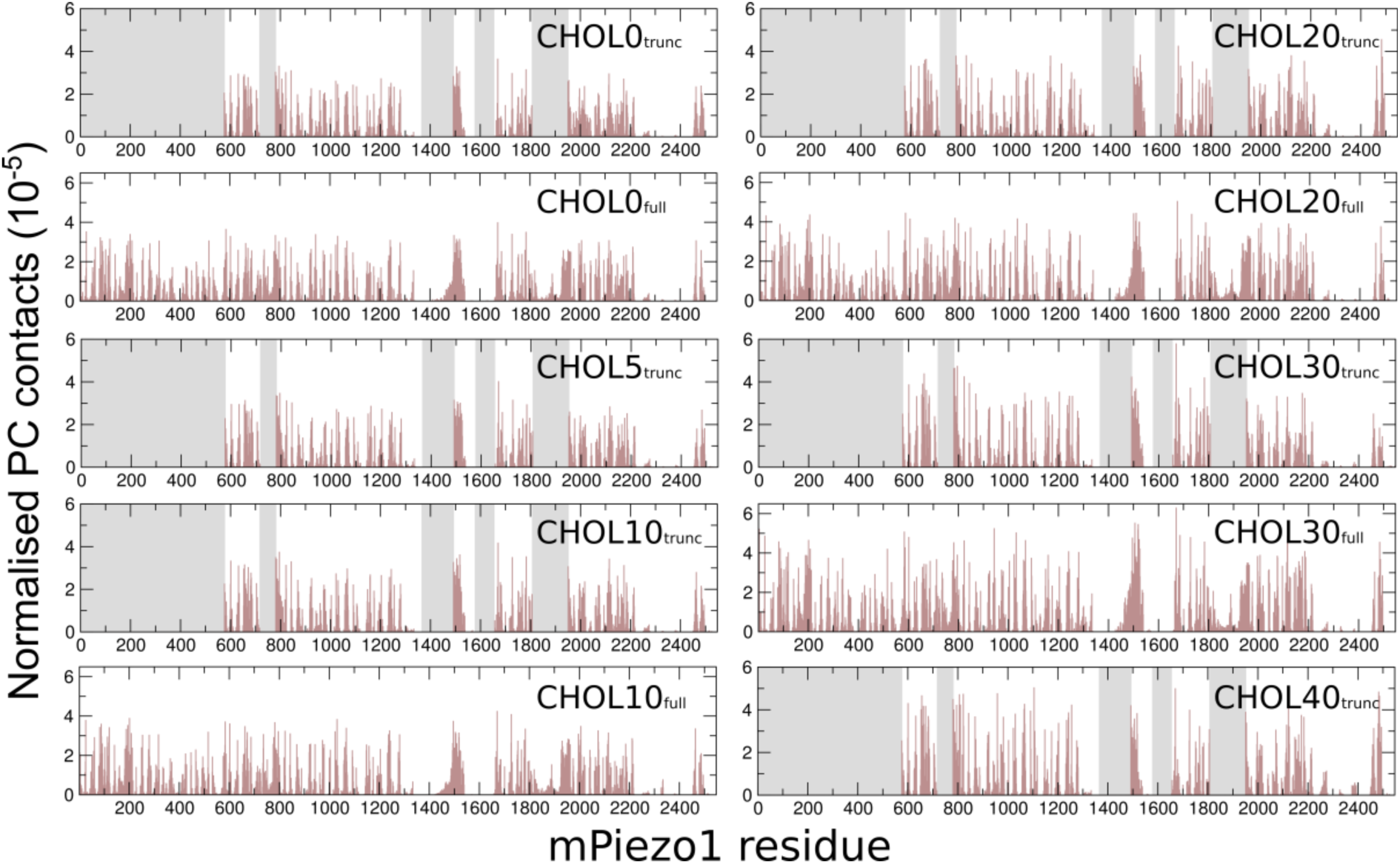
Histograms of Piezo1-POPC contacts in all simulations. Each histogram is labelled with the corresponding simulation. Grey bars represent the residues missing from Piezo1_trunc_.

**Supplementary Figure 8:**
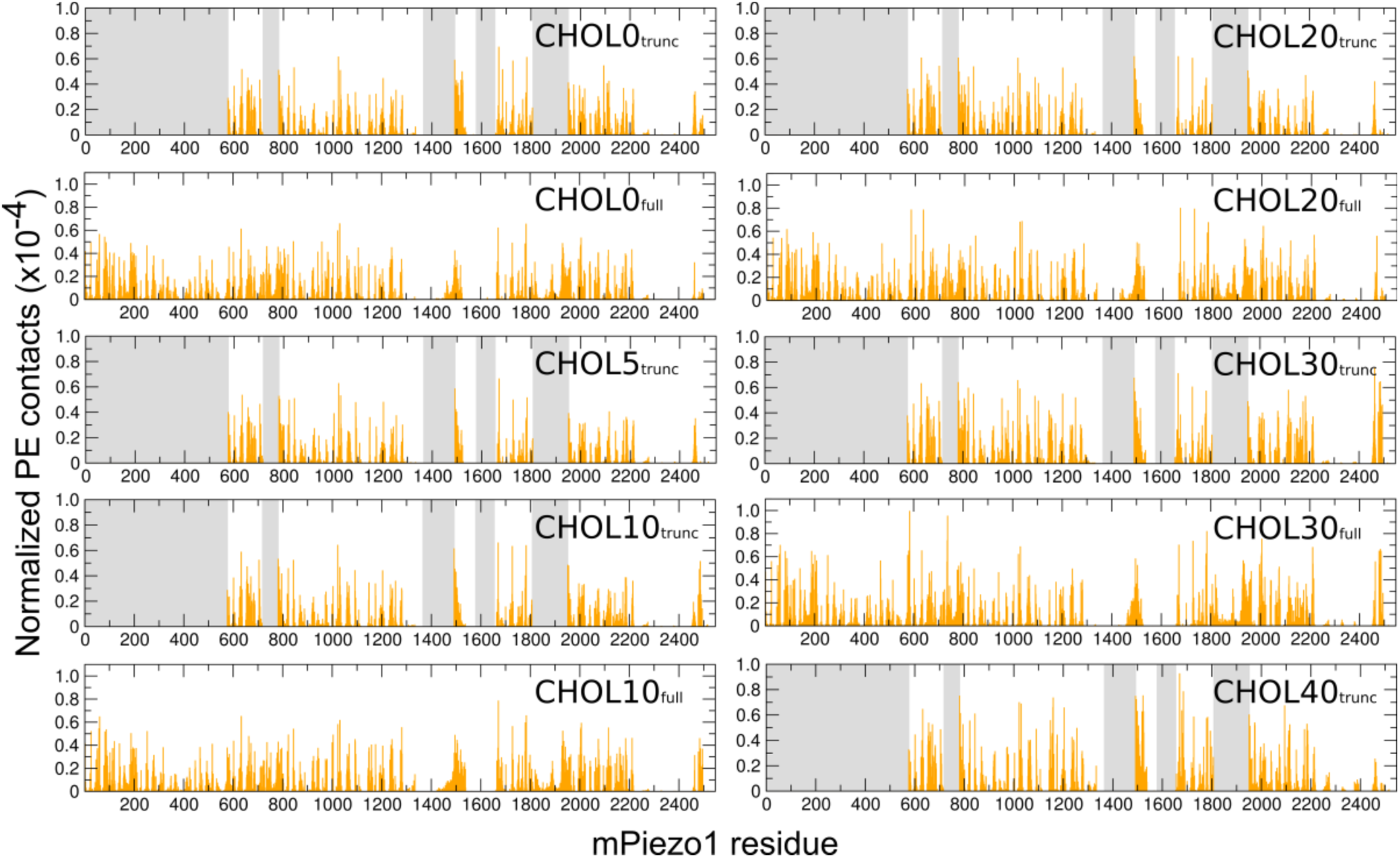
Histograms of Piezo1-POPE contacts in all simulations. Each histogram is labelled with the corresponding simulation. Grey bars represent the residues missing from Piezo1_trunc_.

**Supplementary Figure 9:**
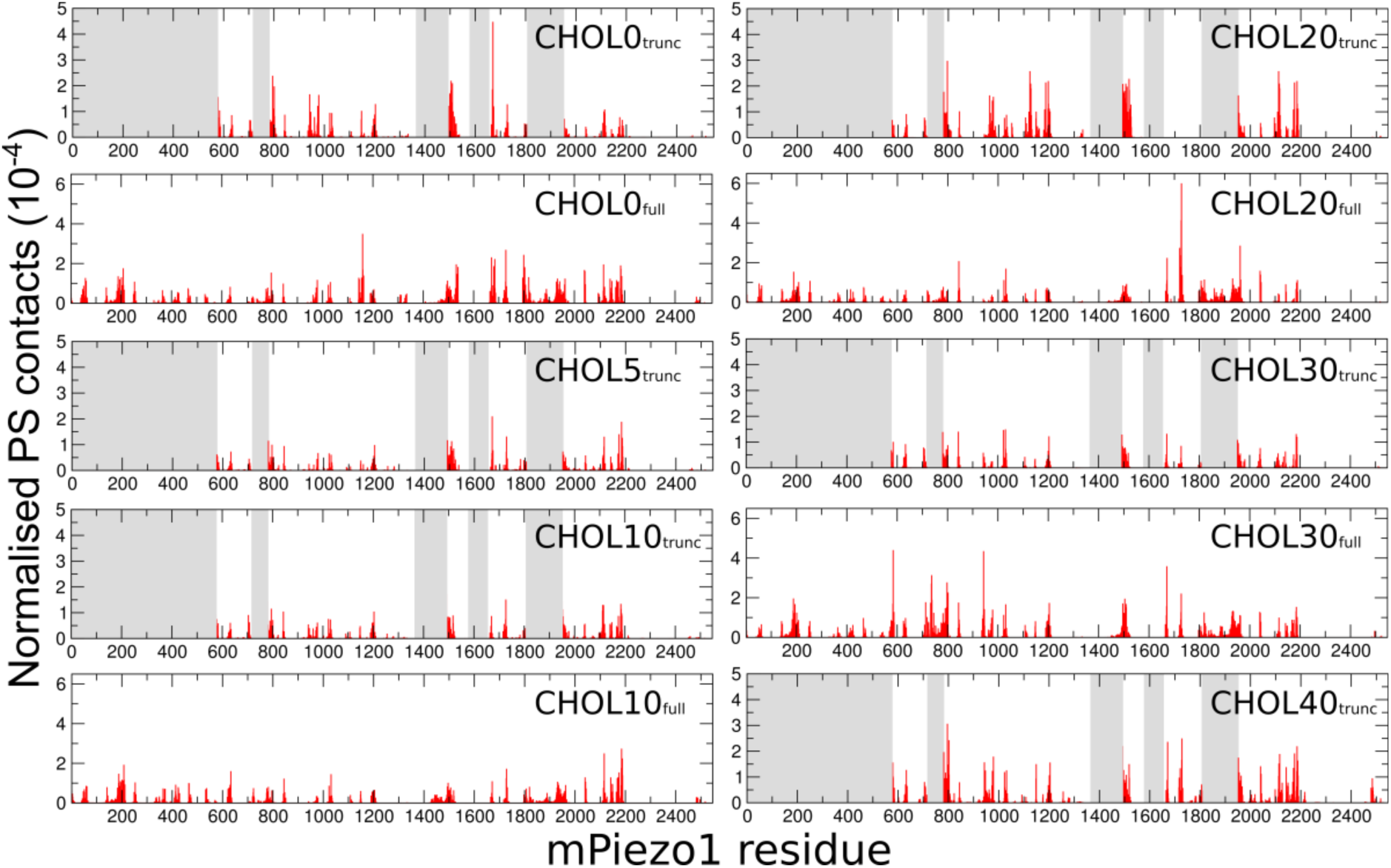
Histograms of Piezo1-POPS contacts in all simulations. Each histogram is labelled with the corresponding simulation. Grey bars represent the residues missing from Piezo1_trunc_.

**Supplementary Figure 10:**
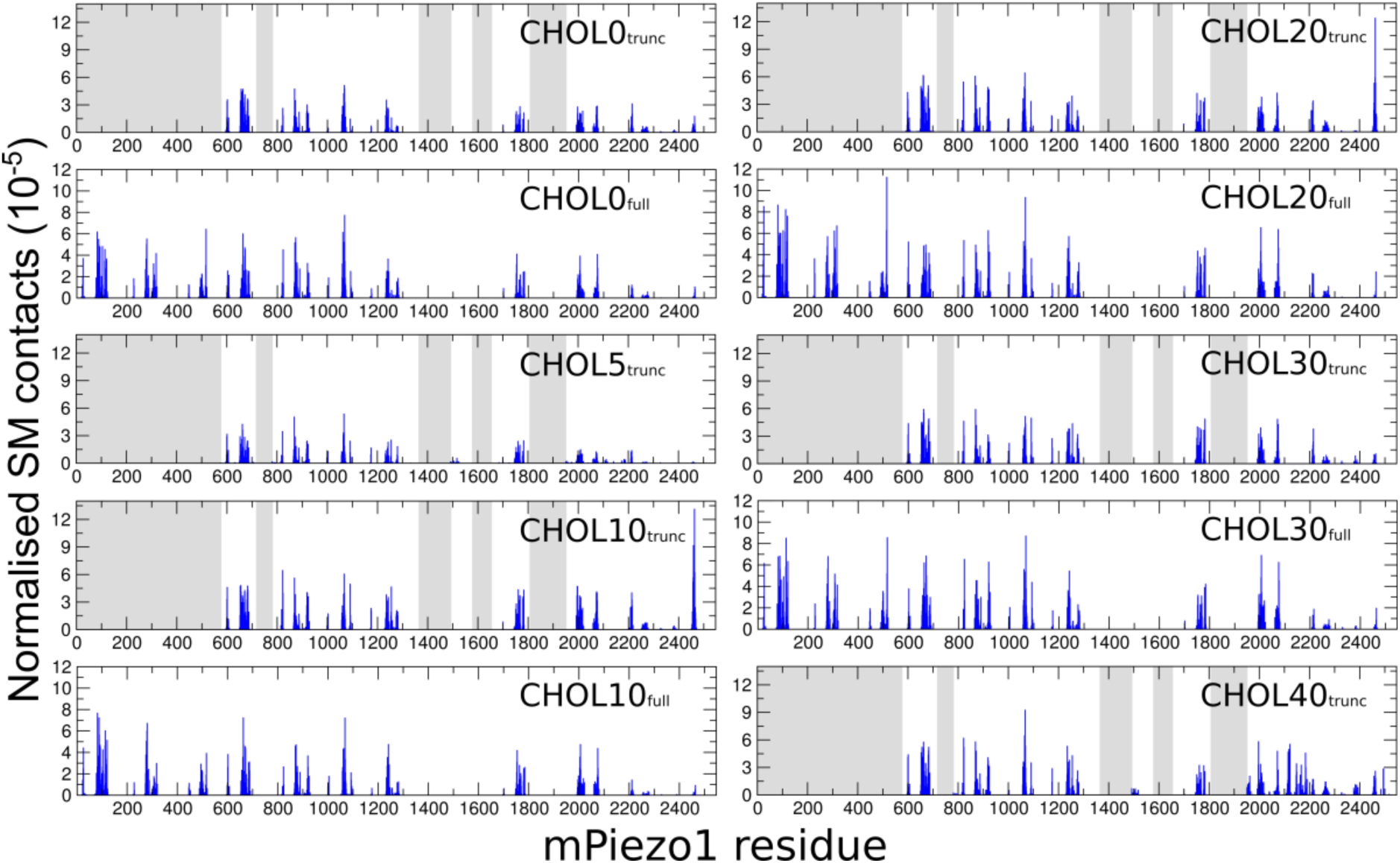
Histograms of Piezo1-SM contacts in all simulations. Each histogram is labelled with the corresponding simulation. Grey bars represent the residues missing from Piezo1_trunc_.

